# Novel genetic variants associated with brain functional networks in 18,445 adults from the UK Biobank

**DOI:** 10.1101/2020.09.17.268029

**Authors:** Heidi Foo, Anbupalam Thalamuthu, Jiyang Jiang, Forrest C. Koch, Karen A. Mather, Wei Wen, Perminder S. Sachdev

## Abstract

This is the first study investigating the genetics of weighted functional brain network graph theory measures from 18,445 participants of the UK Biobank (44-80 years). The eighteen measures studied showed low heritability (mean h^2^_SNP_ =0.12) and were highly genetically correlated. Genome-wide association studies for these measures observed 14 significant variants associated with strength of somatomotor and limbic networks. These intergenic variants were located near the *PAX8* gene on chromosome 2. Gene-based analyses identified five significantly associated genes for five of the network measures, which have been implicated in sleep duration, neuronal differentiation/development, cancer, and susceptibility to neurodegenerative diseases. Genetic correlations with other traits were examined and significant correlations were observed with sleep measures and psychiatric symptoms. Further analysis found that somatomotor network strength was phenotypically associated with sleep duration and insomnia. Single nucleotide polymorphism (SNP) and gene level associations with functional network measures were identified, which may help uncover novel biological pathways relevant to human brain functional network integrity and diseases that affect it.

---

During aging, the human brain undergoes functional changes which affect the integration of information within and between functional brain networks, and these have been shown to be associated with behavioral changes ^1^. By modelling large-scale brain networks using graph theory, defined by a collection of nodes (brain regions) and edges (magnitude of temporal correlation in functional magnetic resonance imaging (fMRI) activity between two brain regions)^2,3^, it is possible to investigate aging-related topological changes. Previous functional graph theory studies have mainly considered edge presence or absence represented as a binary variable^4-6^, which precludes information about variations in connectivity weights between networks. Given that connection weights exhibit high heterogeneity over several orders of magnitude^7^, studying weighted brain networks may provide greater insights into their underlying hierarchy and organizational principles.

Genetics may play an important role in influencing changes in the functional topology of the aging brain. Graph theoretical brain functional measures are reported to be heritable^8,9^. However, to date, there has not been any population-based study investigating the genetic contribution to functional connectivity using graph theory measures. Studying the genetic architecture of graph theory brain functional measures has important implications^10^ - firstly, it can identify genes associated with network topology; and secondly, it may provide insights into the underlying biological mechanisms of macroscopic network topology in aging and how it alters during disease states.

Here, we address the question of how genetics influences the integrity of functional connectivity as measured by graph theory measures in resting-state fMRI (rs-fMRI) data. We assessed graph theory measures, including global efficiency, characteristic path length, Louvain modularity, transitivity, local efficiency and strength of default, dorsal attention, frontoparietal, limbic, salience, somatomotor, and visual networks, which are typically examined and found to change in aging ^6^ as well as multiple neuropathological processes^11-13^. A UK Biobank sample comprising 18,445 participants of British ancestry was used in this study. We first estimated single nucleotide polymorphism (SNP) heritability (h^2^). Subsequently, genome-wide association studies (GWAS) were performed to identify genetic variants associated with each graph theory measure. Gene-based association analysis was carried out to uncover gene-level associations, and the functional consequences of the significant genetic variants were explored.

## Results

### Demographics and graph theory measures

After imaging and genetic pre-processing and quality control, 2,153 UK Biobank participants were excluded resulting in a final sample of 18,445 participants of British ancestry. There were 9,773 females and 8,672 males with a mean age (sd) of 62.47 (7.47). Eighteen weighted graph theory measures were derived by parcellating the rs-fMRI data using the Schaefer atlas^14^, which is a fine-grained parcellation scheme based on Yeo-7 network. These 18 measures include: global efficiency and characteristic path length (network integration); modularity, transitivity, and local efficiency of 7 networks (network segregation); and strength of 7 networks. Supplementary Table 1 defines each of the measures and provides evidence of their association with aging and neuropathological diseases. Supplementary Table 2 shows the mean and standard deviation of the demographics and graph theory measures. Fig. 1 shows the brain network reconstruction using rs-fMRI data.

**Fig. 1.**
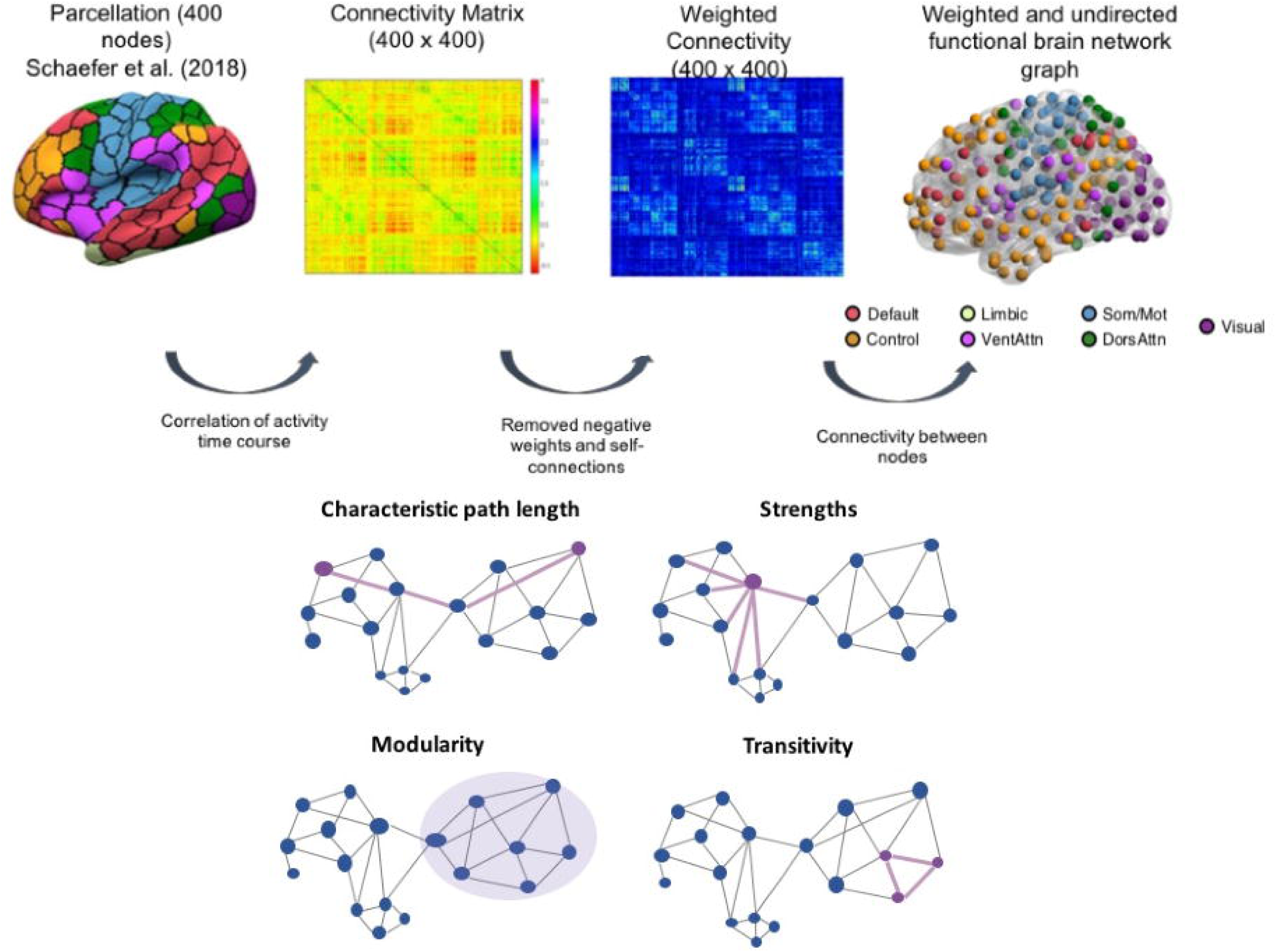
Schematic representation of brain network construction using graph theory analysis. After pre-processing, the brain was divided into different parcels using the Schaefer et al. (2018) parcellation scheme. Subsequently, activity time course was extracted from each region to create the correlation matrix. We used the correlation matrix and removed all the self-connected and negative weights to derive a corresponding weighted undirected brain network matrix and functional brain network graph. Lastly, we used the network matrix to calculate the sets of topological graph theory measures.

Phenotypic correlations between the measures were examined (Supplementary Fig. 1). High correlations were observed between the following measures: characteristic path length and modularity were negatively correlated with all other graph theory measures (r = - 0.661 to −0.989); global efficiency and transitivity were positively correlated with local efficiency and strength of all the networks (r = 0.753 to 0.952); local efficiencies of networks were positively correlated with each other (r = 0.784 to 0.902); and strengths of networks were positively correlated with each other (r = 0.772 to 0.927).

### SNP heritability estimates and genetic correlations

SNP heritability, h^2^_SNP_, was estimated using the proportion of variance in each graph theory measure that is explained by GWAS SNPs. All graph theory measures, except for strength of the visual network, were significantly heritable (*p* < 0.05), ranging from h^2^_SNP_ = 0.07 for local efficiency of visual network to h^2^_SNP_ = 0.17 for the strength of limbic network, with a mean h^2^_SNP_ of 0.12 (Supplementary Table 3; Fig. 2A). Notably, higher h^2^_SNP_ estimates (0.11-0.17) were observed for global efficiency, characteristic path length, transitivity, and strength of limbic, somatomotor, default, salience, and frontoparietal networks compared to the other measures. In addition, genetic and environmental correlations were examined. Strong genetic correlations between the network measures were observed. All measures were positively associated with each other with the exception of Louvain modularity and characteristic path length being negatively correlated with all other measures (Supplementary Table 4; Fig. 2B).

**Fig. 2.**
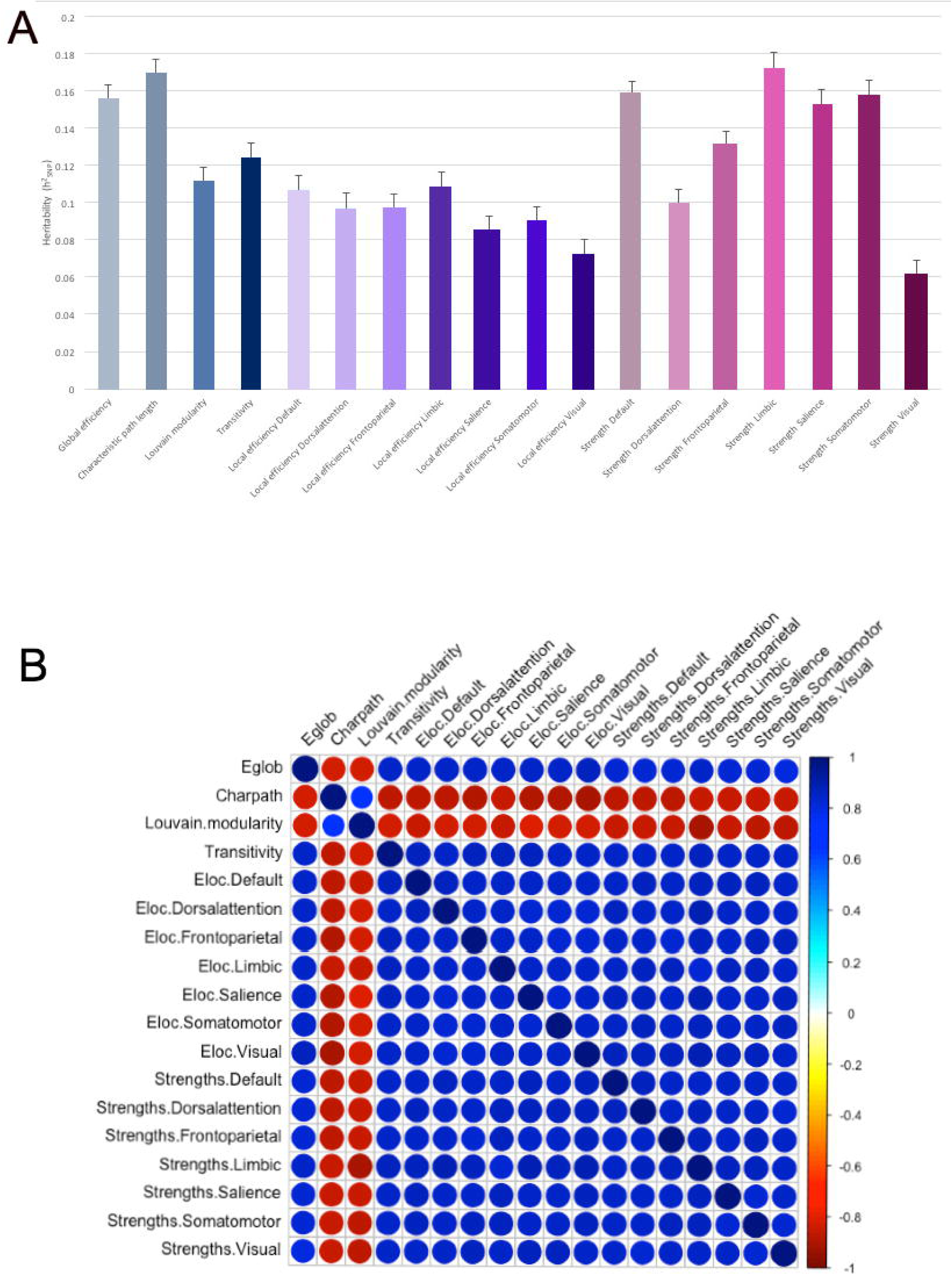
Genetic and environmental correlations between the 18 weighted graph theory measures. (A) represents the heritability estimates for each of the graph theory measures. Gray represents the network integration as characterized by global efficiency and characteristic path length; blue represents network segregation as characterized by Louvain modularity and transitivity; purple represents networks of local efficiency; and pink represents strength of the networks. (B) Genetic correlations were estimated using LDSC (https://github.com/bulik/ldsc). Strong genetic and environmental correlations between the network measures were observed (see Supplementary Table 4 for more details). Abbreviations: Eglob, global efficiency; Charpath, characteristic path length; Eloc, local efficiency.

### Genome-wide association study

GWAS of the rs-fMRI data for each individual graph theory measure (n=18) were carried out using an additive genetic model adjusted for age, age^2^, sex, age × sex, age^2^ × sex, head motion from resting-state fMRI, head position, volumetric scaling factor needed to normalize for head size, genotyping array and 10 genetic principal components.

At the genome-wide significance level of *p* < 5 × 10^−8^ (unadjusted for the number of measures assessed), there were 31 SNPs significantly associated with nine of the 18 graph theory measures namely, global efficiency, characteristic path length, Louvain modularity, local efficiency of default and somatomotor networks, strength of default, limbic, salience, and somatomotor networks (Supplementary Table 5). Supplementary Fig. 2 shows the Manhattan and quantile-quantile plots of all the 18 graph theory measures. However, after adjusting for the number of independent tests using this method ^15^ (n = 6 tests for this study; *p*-threshold = 5 × 10^−8^/6 = 8.33 × 10^−9^), only fourteen variants from a single locus remained significant with strength of the somatomotor network (lead SNP: rs12616641), and one of which also was significant for strength of the limbic network (Table 1; Fig. 3A & 3B). All SNPs were located in an intergenic region near *PAX8* (Paired box gene 8) on chromosome 2 (BP 114065390 - 114092549) (Fig. 3C).

**Table 1.** GWAS genome-wide significant results for brain functional network measures.

| SNP and measures | Chr | Position | Function | Nearest gene | A1/A2 | EAF | $\beta$ | SE | *p* |
| --- | --- | --- | --- | --- | --- | --- | --- | --- | --- |
| **Strength Limbic** |  |  |  |  |  |  |  |  |  |
| rs62158161 | 2 | 114065572 | Intergenic | *PAX8* | C/T | 0.749 | 0.069 | 0.012 | 4.70E-09 |
| **Strength Somatomotor** |  |  |  |  |  |  |  |  |  |
| rs62158160 | 2 | 114065390 | Intergenic | *PAX8* | C/T | 0.742 | 0.066 | 0.011 | 6.90E-09 |
| rs62158161 | 2 | 114065572 | Intergenic | *PAX8* | C/T | 0.749 | 0.069 | 0.012 | 2.40E-09 |
| rs62158166 | 2 | 114077218 | Intergenic | *PAX8* | G/C | 0.773 | 0.073 | 0.012 | 1.20E-09 |
| rs62158168 | 2 | 114078381 | Intergenic | *PAX8* | C/G | 0.773 | 0.073 | 0.012 | 1.20E-09 |
| rs12616641* | 2 | 114079248 | Intergenic | *PAX8* | C/A | 0.774 | 0.073 | 0.012 | 9.90E-10 |
| rs62158169 | 2 | 114081827 | Intergenic | *PAX8* | C/T | 0.784 | 0.072 | 0.012 | 3.50E-09 |
| rs62158170 | 2 | 114082175 | Intergenic | *PAX8* | A/G | 0.784 | 0.073 | 0.012 | 2.60E-09 |
| rs199993536 | 2 | 114082628 | Intergenic | *PAX8* | T/A | 0.783 | 0.072 | 0.012 | 3.10E-09 |
| rs6737318 | 2 | 114083120 | Intergenic | *PAX8* | A/G | 0.779 | 0.070 | 0.012 | 6.20E-09 |
| rs62158206 | 2 | 114084596 | Intergenic | *PAX8* | T/C | 0.779 | 0.071 | 0.012 | 4.10E-09 |
| rs7556815 | 2 | 114085785 | Intergenic | *PAX8* | G/A | 0.780 | 0.071 | 0.012 | 4.70E-09 |
| rs2863957 | 2 | 114089551 | Intergenic | *PAX8* | C/A | 0.779 | 0.071 | 0.012 | 4.10E-09 |
| rs1823125 | 2 | 114090412 | Intergenic | *PAX8* | A/G | 0.779 | 0.071 | 0.012 | 3.30E-09 |
| rs60873293 | 2 | 114092549 | Intergenic | *PAX8* | G/T | 0.779 | 0.070 | 0.012 | 5.70E-09 |

Linear regression models were adjusted for age, age<sup>2</sup>, sex, age × sex, age<sup>2</sup> × sex, head motion from resting-state fMRI, head position, volumetric scaling factor needed to normalize for head size, and 10 genetic principal components. P values are two-tailed. Only results that survived Bonferroni correction ( $p < 5 \times 10^{-8}/6$ ) were reported here.
\*rs12616641 was the lead SNP.
Abbreviations: SNP, single nucleotide polymorphism; Chr, chromosome; A1, coded allele; A2, non-coded allele; EAF, effect allele frequency; $\beta$ , beta; SE, standard error

**Fig. 3.**
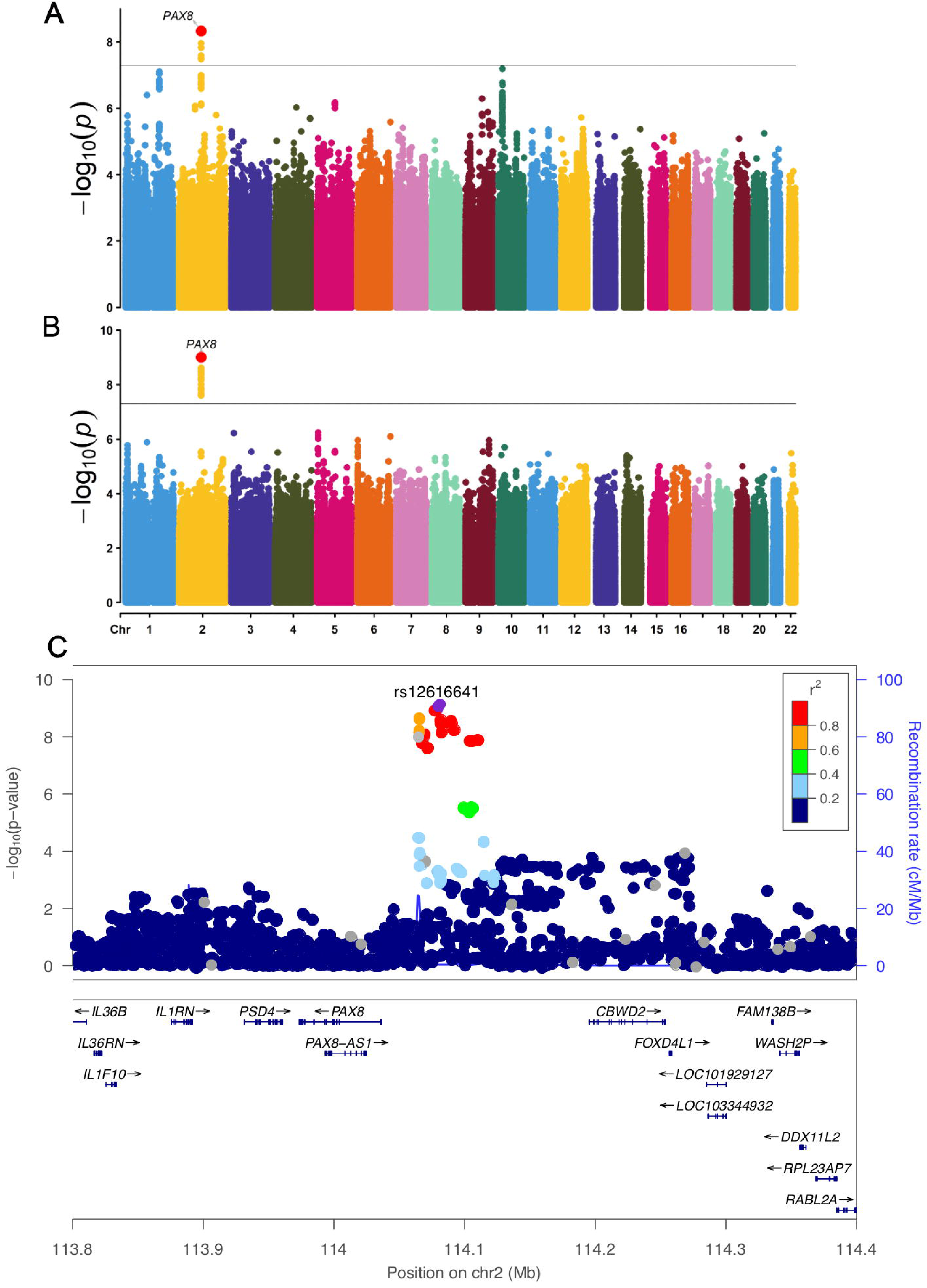
GWAS Manhattan Plots for strength of limbic and somatomotor networks and the locus zoom plot for the identified chromosome 2 region. (A) represents the Manhattan plot for strength of limbic network; (B) represents the Manhattan plot for strength of somatomotor network. For each of the Manhattan plots, each point represents a single genetic variant plotted according to its genomic position (x-axis) and its association with the relevant graph theory measure is shown by the corresponding –log_10_(P) values on the y-axis. Linear regression models were adjusted for age, age^2^, sex, age × sex, age^2^ × sex, head motion from resting-state fMRI, head position, volumetric scaling factor needed to normalize for head size, genotyping array and 10 genetic principal components. The black solid line represents the classical GWAS significance threshold of *p* < 5 × 10^−8^. (C) The Locus zoom plot showing the chromosome 2 locus significantly associated with both the strength of somatomotor and limbic networks. Rs12616641 is the lead SNP.

### Multivariate association test

Since all network measures were highly correlated, to increase power a multivariate analysis using GWAS summary statistics from multiple network measures was performed. Based on their network properties, pooled summary statistics as implemented in metaUSAT^16^ were obtained for (i) global efficiency and characteristic path length; (ii) modularity and transitivity; (iii) local efficiency of all networks; and (iv) strength across all networks. Multivariate analysis of the combined strength network measures yielded 31 significant associations at *p* < 1.25 × 10^−8^ (adjusted for four tests). Out of this, 23 were found in GWAS of somatomotor strength measure reported in supplementary table 1 and the remaining 8 SNPs were found for combined strength measure (Table 2). No additional hits were found in the multivariate analysis of the other three grouped measures.

**Table 2.**
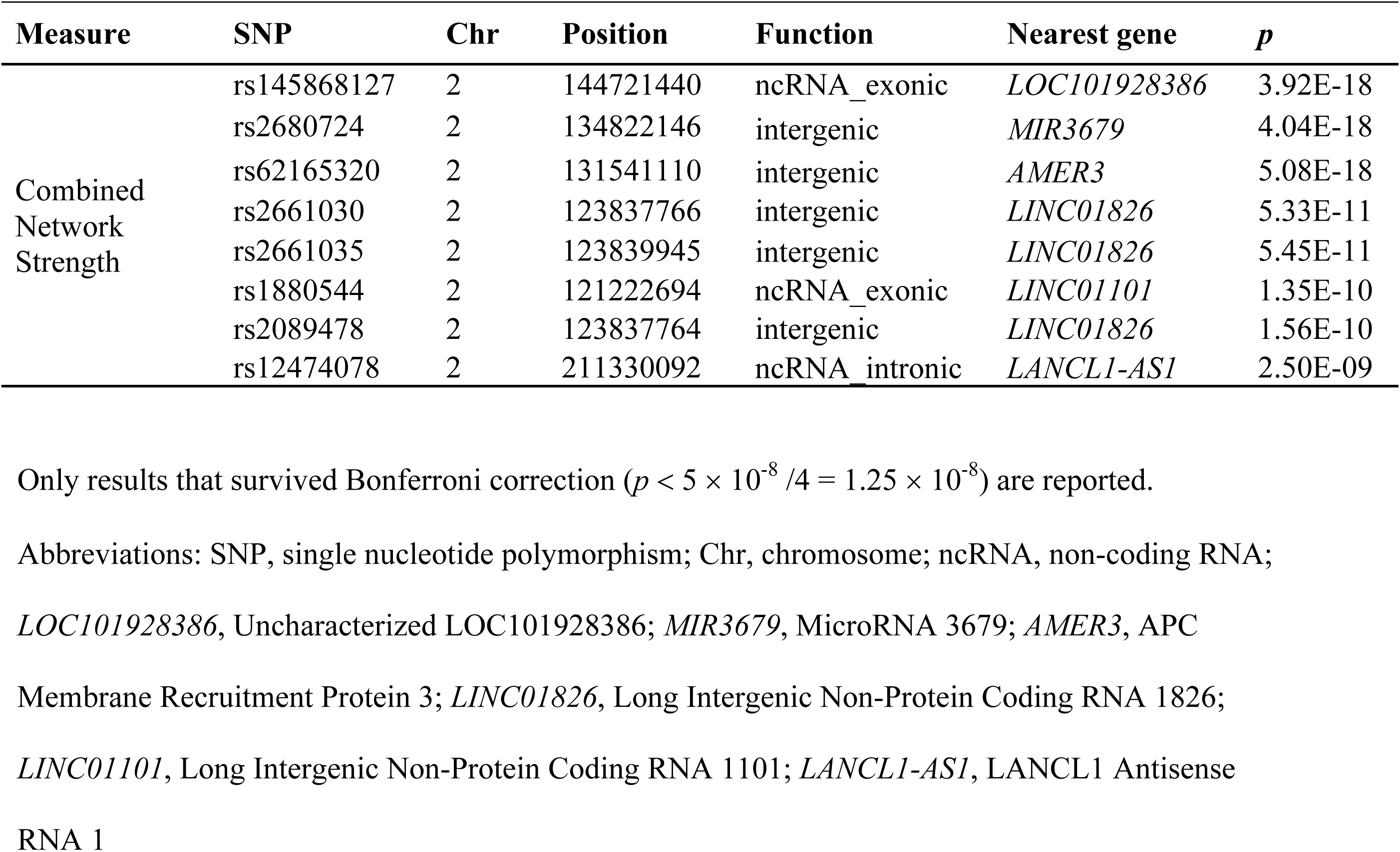
Multivariate SNP-based analyses for combined network strength measure.

### Gene-based association analysis

Gene-based association analysis was performed using the software MAGMA^17^ (Table 3). Our results showed that five genes – *SLC25A33* (Solute Carrier Family 25 Member 33), *TMEM201* (Transmembrane Protein 201), *ZEB1* (Zinc Finger E-Box Binding Homeobox 1), *SH2B3* (SH2B Adaptor Protein 3), and *ATXN2* (Ataxin 2) – were associated with global efficiency, characteristic path length, and strength of default, dorsal attention, and somatomotor networks after adjusting for the number of independent tests (n = 6) and number of genes (n = 18,319) i.e. *p*-threshold < 0.05/(6 × 18319) = 4.56 × 10^−7^. Supplementary Table 6 describes the genes associated with the graph theory measures.

**Table 3.** Gene-based association analysis identified five significant gene-level associations for brain functional network measures.

| Gene | Graph theory measures | Chr | Position | NSNPs | P |
| --- | --- | --- | --- | --- | --- |
| *SLC25A33* | Global efficiency | 1 | 9599528 | 69 | 5.64E-08 |
|  | Characteristic path length | 1 | 9599528 | 69 | 5.23E-08 |
| *TMEM201* | Global efficiency | 1 | 9648932 | 43 | 1.83E-08 |
|  | Characteristic path length | 1 | 9648932 | 43 | 1.48E-08 |
|  | Strength Dorsal attention | 1 | 9648932 | 43 | 1.36E-07 |
|  | Strength Somatomotor | 1 | 9648932 | 43 | 2.09E-07 |
| *ZEB1* | Global efficiency | 10 | 31607825 | 323 | 3.27E-07 |
|  | Characteristic path length | 10 | 31607825 | 323 | 2.29E-07 |
| *SH2B3* | Strength Default | 12 | 111843720 | 50 | 2.95E-07 |
| *ATXN2* | Strength Default | 12 | 111890018 | 149 | 2.48E-07 |

Gene-based association analysis was performed via MAGMA<sup>17</sup>, which uses 1000G reference panel for calculation of LD between the SNPs and gene coordinates based on NCBI build 37. Only genes that passed the Bonferroni correction ( $p < 4.56 \times 10^{-7}$ ) are reported here.
Abbreviations: Chr, chromosome; NSNP, number of single nucleotide polymorphisms
*SLC25A33*, Solute Carrier Family 25 Member 33; *TMEM201*, Transmembrane Protein 201; *ZEB1*,
Zinc Finger E-Box Binding Homeobox 1; *SH2B3*, SH2B Adaptor Protein 3; *ATXN2*, Ataxin 2

### Functional annotations

We assessed the potential functions of the 39 significant SNPs from the GWAS (n=31) and multivariate association tests (n=8) in SNPnexus^18^. The results showed that many of the network associated variants were associated with other phenotypes including sleep patterns, psychiatric disorders, coronary artery disease, cholesterol, and blood pressure (Tables S7 and S8). SNPnexus did not annotate any of the variants as eQTLs.

Gene expression enrichment analysis of the list of significant genes in our results was subsequently performed using the Functional Mapping and Annotation (FUMA) platform^19^. They were expressed in the brain but also across other tissue types (Supplementary Fig. 3). Moreover, there was no enrichment in brain tissue types (Supplementary Table 9).

### Genetic correlations with other traits

Given that the most robust GWAS result was observed for the strength of somatomotor network, we examined its genetic correlations with other traits via linkage disequilibrium (LD) score regression (LDSC)^20^ (Supplementary Table 10). Results showed that strength of somatomotor network was genetically correlated with other traits, such as depressive symptoms and subjective well-being (unadjusted *p* ≤ 0.05), as well as nearly significant correlations with sleeping behaviours, cognitive and other psychiatric symptoms.

### Correlations with other associated traits

As the SNPs in our study have been associated with sleep and insomnia in previous GWAS studies (Supplementary Table 7), we explored the phenotypic correlations between the graph theory and sleep-related measures in the UK Biobank data (Tables S11 and S12). Self-reported sleep duration and insomnia were highly associated with the graph theory measures. Strength of somatomotor network showed the most significant association with sleep duration (*p* = 8.33 × 10^−11^) and insomnia (*p* = 0.0019).

## Discussion

Functional graph theory measures reflect the underlying functional topography of the brain. To date, this is the first study investigating the genetics of weighted functional graph theory measures. We present h^2^_SNP_ estimates and results from GWAS of graph theory measures using resting-state fMRI data from 18,445 UK Biobank participants. We identified significant SNPs and gene associations that survived multiple correction testing with six of the 18 graph theory measures including global efficiency, characteristic path length, and strength of default, dorsal attention, limbic, and somatomotor networks. The novel contributions of this paper are the identification of new genetic associations at the variant, locus, and gene levels, providing insights into the genetic architecture of graph theory metrics using resting-state fMRI data.

We found low h^2^_SNP_ across all the graph theory measures. This is in contrast to previous classical twin study design studies that have shown moderate to high heritability of 0.52 to 0.64 for global efficiency, 0.47 to 0.61 for mean clustering coefficient, and 0.38 to 0.59 for modularity^8^. One of the plausible explanations is that h^2^_SNP_ typically provides smaller heritability estimates due to uncaptured rare genetic variants compared to those provided by the classic twin study design^21^. High genetic correlations observed between graph theory measures implies that there may be common variants contributing to these measures.

The GWAS results found that strength of limbic and somatomotor networks were associated with SNPs located in an intergenic region near *PAX8* (Paired box gene 8) on chromosome 2, which is the closest gene to the top SNP. The *PAX* gene family encodes transcription factors which are essential during development and tissue homeostasis^22^. Specifically, *PAX8* protein is considered as a master regulator for key cellular processes in DNA repair, replication, and metabolism^23^. It has also been shown to regulate several genes involved in the production of thyroid hormone^24^, essential for brain development and function such as neuronal differentiation, synaptogenesis, and dendritic proliferation^25,26^. A previous study has also linked reductions in intrinsic functional connectivity in the somatomotor network to participants with subclinical hypothyroidism compared to controls^27^. Interestingly, studies have also found subclinical thyroid dysfunction to be associated with sleep quality^28,29^. Considering that we also observed phenotypic correlations between sleep duration, insomnia and graph theory measures, it is possible that *PAX8* variants may play a role in the regulation of genes associated with functional brain network properties and sleep.

Results from multivariate SNP-based analyses from combined network strength measures found associations with SNPs on chromosome 2, which were associated with inflammation and oncogenesis. The majority of these SNPs were located in intergenic regions close to non-coding RNA genes. *LINC01826* (long intergenic non-protein coding RNA 1826) has been associated with inflammatory responses of the vascular endothelial cells, which are important in the development of cardio-cerebrovascular diseases^30^. *MIR3679* (microRNA 3679) is a short non-coding RNA involved in post-transcriptional regulation of gene expression, which has been postulated to function as a tumor suppressor due to lower levels observed in patients with diffuse glioma than controls^31^. *LINC01101* (long intergenic non-protein coding RNA 1101), on the other hand, has been down-regulated in cervical cancer^32^. The identified genes may contribute to understanding the relationship between strength of brain networks and disease.

Gene-based association analysis showed associations with genes involved in neuronal differentiation/development, cancer, and susceptibility to neurodegenerative diseases. *ZEB1* has been implicated in neuronal glioblastoma^33^ whereas *SLC25A33* has been associated with insulin/insulin-like growth factor 1 (IGF-1) necessary for metabolism, cell growth, and survival^34^ and showed higher expression in transformed fibroblasts and cancer cell lines compared to non-transformed cells^34,35^. *TMEM201* mice, which undergo accelerated senescence, exhibited an early onset age-related decline in antibody response and have a higher rate of mortality^36^. *ATXN2* belongs to a class of genes associated with microsatellite-expansion diseases, where an interrupted CAG repeat expansion has been associated with brain-related diseases including spinocerebellar ataxia type 2 (SCA2), frontotemporal lobar degeneration (FTLD), and amyotrophic lateral sclerosis (ALS)^37,38^. A neighbouring gene *SH2B3* to *ATXN2* has also been implicated in increased ALS risk^38^. Consistent with previous studies that identified functional graph theory measures that were associated with the most common FTLD (behavioral variant of frontotemporal dementia)^39,40^, we observed similar graph theory measures to be associated with the *ATXN2* gene. This implies that *ATXN2* may be involved in the relationship between the disruption of brain networks and neurological/neuromuscular disorders.

The strengths of this study include the well characterized sample and its large sample size, and uniform MRI methods. The results, however, should be interpreted with caution. While using weighted undirected matrix circumvents issues surrounding filtering/thresholding the connectivity matrix to maintain significant edge weights represented in a binary matrix, there are inherent difficulties associated with the interpretation of the results. As brain signals recorded from resting-state fMRI are typically noisy, it is possible that edge weights may be affected by non-neural contributions^41^. Despite this, with careful denoising of the resting-state fMRI data^42,43^ and covarying for motion, it is possible to minimize the noise in the data. In addition, previous studies have suggested that stronger edge weights make greater contributions in the computation of graph metrics than lower weight connections^44,45^. This implies that when evaluating weighted graphs, false positive connections based on lower correlations may have a less disruptive impact on the network topology^46^. Given that the brain is a complex system with hierarchical network structure, studying weighted networks, as was done in this study, may provide a more holistic representation of the brain functional network. Future studies may benefit from investigating the genetic effects between binarized and weighted graph theory metrics. In addition, the lack of replication cohort is another limitation which can be corrected when independent large datasets become available.

In summary, this is the first study to investigate the genetics of weighted functional graph theory measures in a large and well characterized cohort. We observed multiple SNPs and genes associated with weighted graph theory measures, which have been observed to be implicated in sleep duration, neuronal differentiation/development, cancer, and susceptibility to neurodegenerative diseases. Our findings may help in the identification of novel biological pathways relevant to human brain functional network integrity and disease.

## Acknowledgements

H.F. is supported by the University of New South Wales (UNSW) Scientia PhD scholarship. We gratefully acknowledge that this research has been conducted using the UK Biobank Resource (Application number: 45262).

## Author contributions

H.F. devised the study; H.F., and A.T. jointly designed the analyses, analyzed the data, interpreted and wrote the paper with technical support and guidance with data interpretation from K.M., W.W., and P.S.; H.F., A.T., J.Y.J., and F.C.K. contributed to the statistical concepts; and P.S. supervised the work. All authors commented on the manuscript.

## Data and material availability

Access to cohort data is available through application to the UK Biobank repository. Most of the software and codes used in this study are publicly available. All graph theory measures executed using BCT are available at https://sites.google.com/site/bctnet/Home. This toolbox is currently licensed free and uses MATLAB. The specific scripts for each measure can be found at https://github.com/heidifoo/bctgraphtheory. SNP heritability analysis was performed using https://cnsgenomics.com/software/gcta/#Overview. GWAS was derived using https://data.broadinstitute.org/alkesgroup/BOLT-LMM/#x1-310005.2.

## List of Supplementary Materials

Supplementary Figures 1–3

Supplementary Tables 1–12

## Methods

### Study population

Our study was approved by the UK Biobank in December 2018 (Application number: 45262). rs-fMRI data for 20,598 participants with British ancestry was downloaded in March 2019^47^. The imaging assessment took place at three different assessment centres with the majority assessed in Manchester, more recently scans were undertaken at Newcastle and Reading, UK. The UK Biobank study was conducted under approval from the NHS National Research Ethics Service (approval letter dated 17th June 2011, ref. 11/NW/0382), project 10279. All data and materials are available via UK Biobank (http://www.ukbiobank.ac.uk). Only individuals with both genetics and rs-fMRI data were included in this study.

### Image processing

Briefly, the UK Biobank structural T1-weighted MRI scans were acquired on three 3T Siemens Skyra MRI scanners (software platform VD13) at three sites (Reading, Newcastle, and Manchester) using a 32-channel receiving head coil and a 3D MPRAGE protocol (1.0 x 1.0 x 1.0 mm resolution, matrix 208 x 256 x 256, inversion time (TI)/repetition time (TR) = 880/2,000 ms, in-plane acceleration 2). An extensive overview of the data acquisition protocols and image processing carried out on behalf of the UK Biobank can be found elsewhere^48^. Rs-fMRI data previously pre-processed by the UK Biobank ^48^ were used. Briefly, motion correction, intensity normalization, highpass temporal filtering, echo-planar imaging (EPI) unwarping, and gradient distortion correction were performed. Subsequently, structured artefacts were removed by ICA+FIX processing (Independent component analysis followed by FMRIB’s ICA-based X-noiseifier)^49-51^. We removed participants with motion of > 2mm/degrees of translation/rotation. After image quality control and removal of participants with head motion outliers, we excluded 1,626 participants and 18,972 participants remained.

### Graph theory analyses

The Schaefer atlas^14^ was used as it met the following requirements: 1) it integrated both local gradient and global similarity approaches (i.e. Markov Random Field model) for parcellation, which showed greater homogeneity than four other previously published parcellations implying that it will not overestimate local connectivity of the regions; 2) it revealed neurobiologically meaningful features of the brain organization; and 3) it was based of the Yeo 7-parcels atlas^52^ and provided a more fine-grained parcellation, which allowed us to look at an average of the parcels across the 7 networks for local efficiency and nodal strength. 3dNetCorr command from Analysis of Functional Neuroimages (AFNI)^53^ was used to produce network adjacency matrix for each participant. The mean time-series for each region was correlated with the mean time-series for all other regions and extracted for each participant. Partial correlation, *r*, between all pairs of signals was computed to form a 400-by-400 (Schaefer atlas) connectivity matrix, which was then Fisher z-transformed. Self-connections and negative correlations were set to zero. The main network analysis was performed on positive weighted networks. Given that connection weights in brain networks can vary across magnitude, undirected weighted connectivity matrices were used instead of binary connectivity matrices. The higher the weight, the stronger the functional connectivity is between the brain regions^41^.

All graph theory measures were quantified by using the brain connectivity toolbox (BCT)^2^. Global-level measure included global efficiency, characteristic path length, transitivity, and Louvain modularity. A single value is derived for each of these four measures and quantitatively represent the whole brain network. Network level measures, such as local efficiency and strength, were estimated for each node and averaged across the all subjects within each network. Subsequently, we averaged the left-right hemisphere to derive a value for each node and averaged within each network to derive a value for each of the 7 networks.

To assess integration of information, we calculated global efficiency and characteristic path length^2,54^. Global efficiency represents how effectively the information is transmitted at a global level and is the average inverse shortest path length while the latter measures the integrity of the network and how fast and easily information can flow within the network. To assess network segregation, which characterizes the specialized processing of the brain at a local level, we calculated the Louvain modularity, transitivity, and local efficiency indices^2,54^. Louvain modularity is a community detection method, which iteratively transforms the network into a set of communities, each consisting of a group of nodes. Higher modularity values indicate denser within-modular connections but sparer connections between nodes that are in different modules. Transitivity refers to the sum of all the clustering coefficients around each node in the network and is normalized collectively. Local efficiency is a node-specific measure and is defined relative to the sub-graph comprising of the immediate neighbours of a node. Finally, strength (weighted degree) is described as the sum of all neighbouring edge weights^2^. High connectivity strength indicates stronger connectivity between the regions, which provides an estimation of functional importance of each network.

### Genotype data in UK Biobank

Genetic data for approximately 500,000 were available and full details on the genetics data used were described previously^55^. The samples were collected from stored blood samples in the UK Biobank and genotyped either using the UK Bileve or the UK biobank axiom array. Genotyping was performed on 33 batches of ∼4,700 samples by Affymetrix (High Wycombe, UK). Further details on the UK Biobank sample pre-processing are available here http://biobank.ctsu.ox.ac.uk/crystal/refer.cgi?id=155583.

An imputed data set was made available where the UK Biobank interim release was imputed to a reference set that consisted of > 92 million autosomal variants imputed from the Haplotype Reference Consortium (HRC)^56^ and UK10K + 1000 Genomes resources reference panels.. After SNP QC filters (MAF > 0.1% and imputation information score > 0.3), 9926107 SNPs were used in the GWAS analysis.

Further quality control measures were performed. Due to the confounds associated with population structure^57^, only samples repored to have recent British ancestry were used in the GWAS analysis. Outliers, including those with high missingness, relatedness, quality control failure, and gender mismatch, were removed. The final UK Biobank sample, after genotyping quality control and including those with rs-fMRI data, was n = 18,445 participants.

### Sleep behavioral data in UK Biobank

Self-reported sleep data, namely sleep duration and frequency of insomnia, were used. Sleep duration was recorded as the number of reported hours of sleep in every 24 hours and frequency of insomnia was recorded as never/rarely, sometimes, or usually. More details can be found on http://biobank.ndph.ox.ac.uk/showcase/field.cgi?id=1160 and http://biobank.ndph.ox.ac.uk/showcase/field.cgi?id=1200.

### Potential confounds

In accordance with previous GWAS of brain imaging phenotypes in the UK Biobank^21^, we controlled for similar confounding variables in this study. In addition to age at scanning and sex (i.e. age, age^2^, age × sex, age^2^ × sex), covariates relating to imaging parameters and genetic ancestry were also included. These included: head motion from resting-state fMRI, head position, volumetric scaling factor needed to normalize for head size, and the 10 genetic principal components.

### SNP Heritability

Heritability analysis, which is defined as the proportion of observed phenotypic variance explained by additive genetic factors of all common autosomal variants^58^, estimates the relative contribution of genes and the environment on a phenotype. Using BOLT-REML^59^ implemented in BOLT-LMM v2.3^60^, we estimated the heritability accounted by autosomal SNPs among the graph theory measures. BOLT-REML uses multiple component modelling to partition SNP heritability and applies Monte Carlo algorithm for variance component analysis.

### Genome-wide association analysis

BOLT-LMM v2.3^60^ was used to conduct GWAS for the graph theory measures in the UK Biobank sample and adjusted for potential confounds. To correct for multiple hypothesis testing, we estimated the number of independent tests used based on Nyholt et al. method^15^ and derived n = 6 independent tests for this study. The genome-wide significant threshold was set at *p* < (5 × 10^−8^ /6) = 8.33 × 10^−9^. Findings with unadjusted threshold of *p* < 5 × 10^−8^ were also reported in Supplementary Table 5. Quantile and Manhattan plots were also presented for each of the graph theory measure in Supplementary Fig. 2. Manhattan and QQ plots were made using the R-package^61^. Locus Zoom^62^ was used for the visualization of the nearest genes within a ±500-kilobase genomic region for the strength of somatomotor and limbic networks based on the hg19 UCSC Genome Browser assembly (Fig. 3).

### Multivariate association analysis

Multivariate association analysis was conducted using metaUSAT^16^. The method uses summary statistics from individual studies and is suitable for correlated traits. MetaUSAT derives strength from two methods of multivariate association tests (score test and multivariate analysis variance test) as it uses convex linear combination of two test statistics. Data-driven minimum p-value corresponding to the best-linear combination is obtained. The significant threshold was set as *p* < (5 × 10^−8^ /4) = 1.25 × 10^−8^.

### Gene-based association analysis and functional mapping

Gene-based association analysis was performed via MAGMA^17^ (v1.07, https://ctg.cncr.nl/software/magma/). MAGMA uses 1000G reference panel for calculation of LD between the SNPs and gene coordinates based on NCBI build 37. Gene-based association test statistic was derived using the default option, which is the sum of –log(SNP p-value). The Bonferroni correction was used for significance of the gene-based association tests (p-value of the gene-based tests / (number of independent tests × number of genes tested)). In other words, significant threshold was set as *p* < 0.05/(6 × 18319) = 4.56 × 10^−7^.

### Functional annotation

We performed lookups for variants that passed the suggestive GWAS threshold of *p* < 5 × 10^−8^ to investigate the previously reported associations with the other traits. NHGRI-EBI GWAS Catalogue^63^ included previous GWAS publications (Supplementary Table 7) and SNPnexus^18^ included all other publications (Supplementary Table 8).

FUMA^19^ gene2func online platform (version 1.3.4, http://fuma.ctglab.nl/) was used to explore the functional consequences of significant genes. All the genes in the GWAS analysis and the gene-based analysis after nominal significance were used as input (a total of 18 genes). Using the FUMA platform, the GTEx v7 30 general tissue types data set was used for tissue specificity analyses. Differentially expressed gene (DEG) sets were pre-calculated by performing two-tailed t-test for any one of the tissue type against all others. Expression values were normalized (zero-mean) following a log_2_ transformation of expression values. Using the genes as background gene set, 2 × 2 enrichment sets were performed. Genes with *p-*value ≤ 0.05 were after Bonferroni correction were defined as differentially expressed and colored in red in Supplementary Figure 3.

### Genetic correlation estimation with LDSC

LD Hub (v1.9.1, http://ldsc.broadinstitute.org/ldhub/) was used to estimate the genetic correlation between graph theory measures and corresponding traits. Summary statistics were uploaded to LD hub where it calculates the genetic correlations using the LDSC software (v1.0.0, https://github.com/bulik/ldsc).

### Test of associations with graph theory measures

The graph theory measures were normalized using ranked transformed using the rntransform() function in R from GeneABEL package^64^ and age were z-transformed for regression analysis. Regression model similar to GWAS analysis was used to test the association of self-reported sleep duration and frequency of insomnia variables with the graph theory measures. Only results with a two-tailed *p* < 0.05/18 (number of graph theory measures) were considered significant. Data management, derivation of summary statistics and other statistical analyses, and correlation plots were performed using R (V 4.0.0) software^61^.

