## Supplementary Figures for "Novel genetic variants associated with brain functional networks in 18,445 adults from the UK Biobank"

**Contents**

Supplementary Fig. 1 ………………………………………………………………………. 2

Supplementary Fig. 2 ……………………………………………………………………… 3

Supplementary Fig. 3 ……………………………………………………………………… 22

**Supplementary Fig. 1**. ***Correlations between the graph theory measures in the UK Biobank sample (n = 18,445).*** Blue represents positive correlations whereas red represents negative correlations.

**UK Biobank**

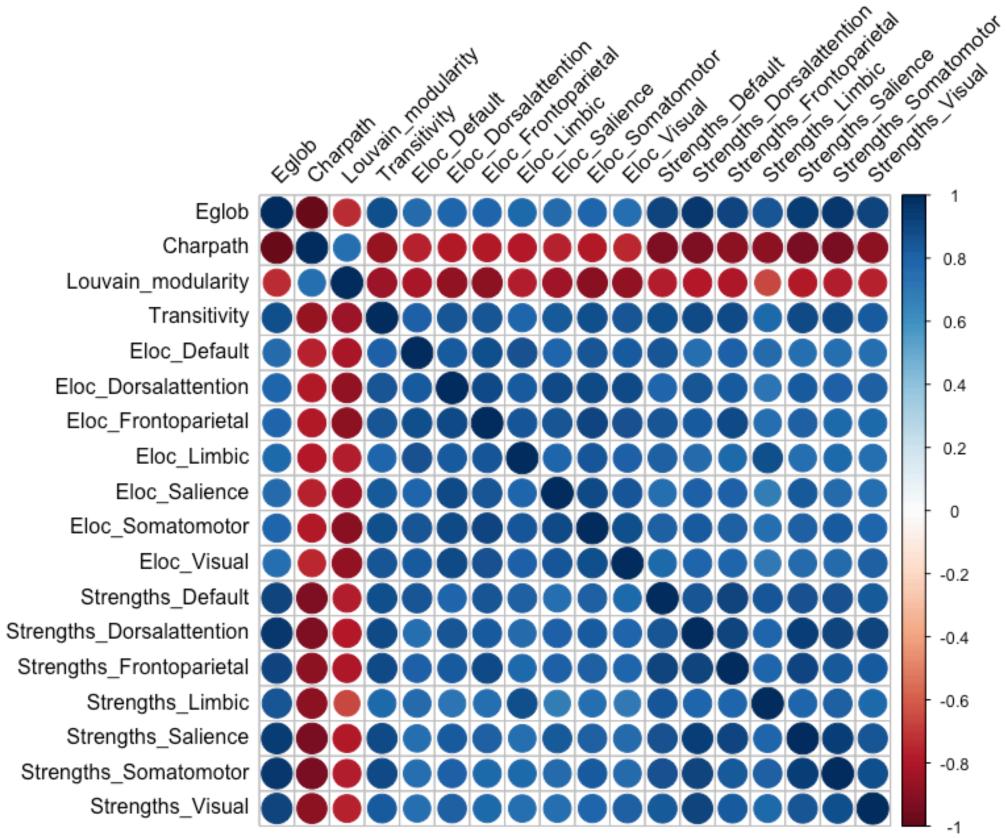


Abbreviations: Eglob, global efficiency; Charpath, characteristic path length; Eloc, local efficiency

**Supplementary Fig. 2.** ***Manhattan and quantile-quantile (QQ) plots in the UK Biobank sample (n = 18,445).*** In the Manhattan plots, each point represents a single genetic variant plotted according to its genomic position (x-axis) and its $-\log_{10}(P)$ for two-tailed associations with graph theory measures (y-axis). In QQ-plots, the line represents the expected null distribution and Lambda inflation factors ($\lambda$) are provided in each plot. Linear regression models were adjusted for age, $age^2$, sex, age × sex, $age^2$ × sex, head motion from resting-state fMRI, head position, volumetric scaling factor needed to normalize for head size, genotyping array, the 10 genetic principal components. The black solid line represents the classical GWAS significance threshold of $p < 5 \times 10^{-8}$. Genomic inflation factor (lambda gc) is between 1.013 to 1.046.

A. Global efficiency

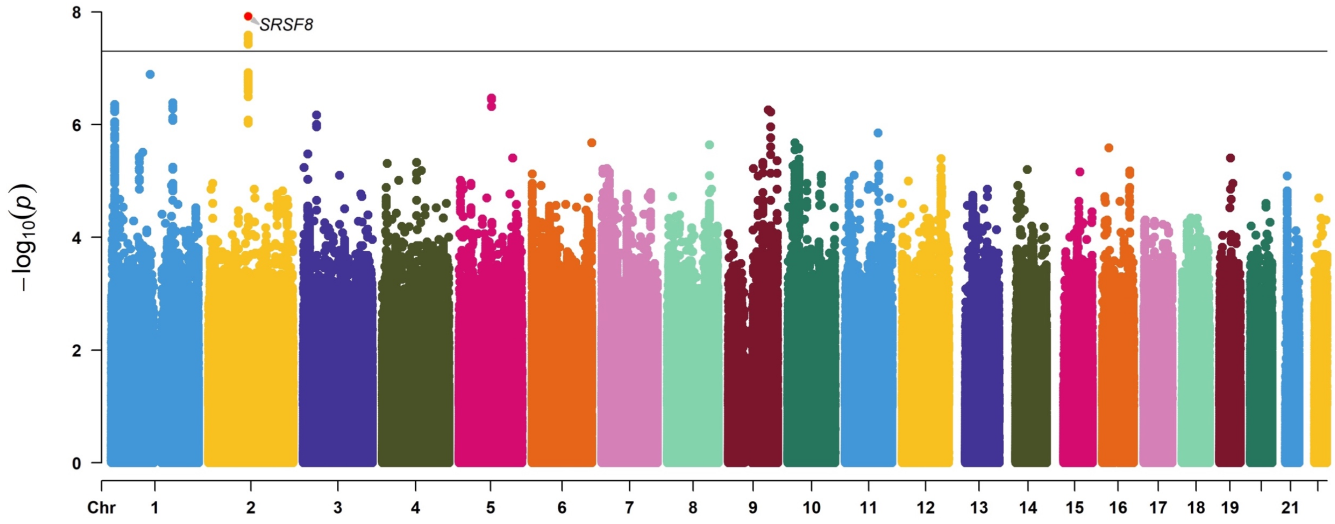


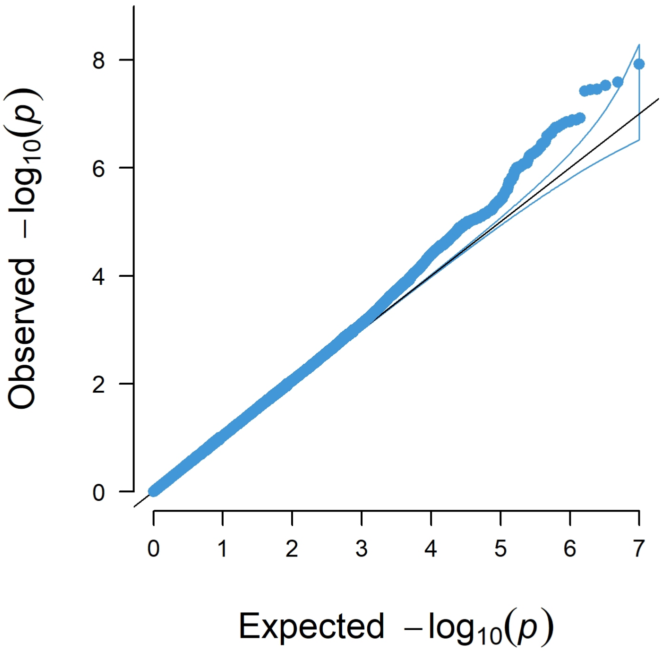


**Lambda gc = 1.043**

B. Characteristic path length

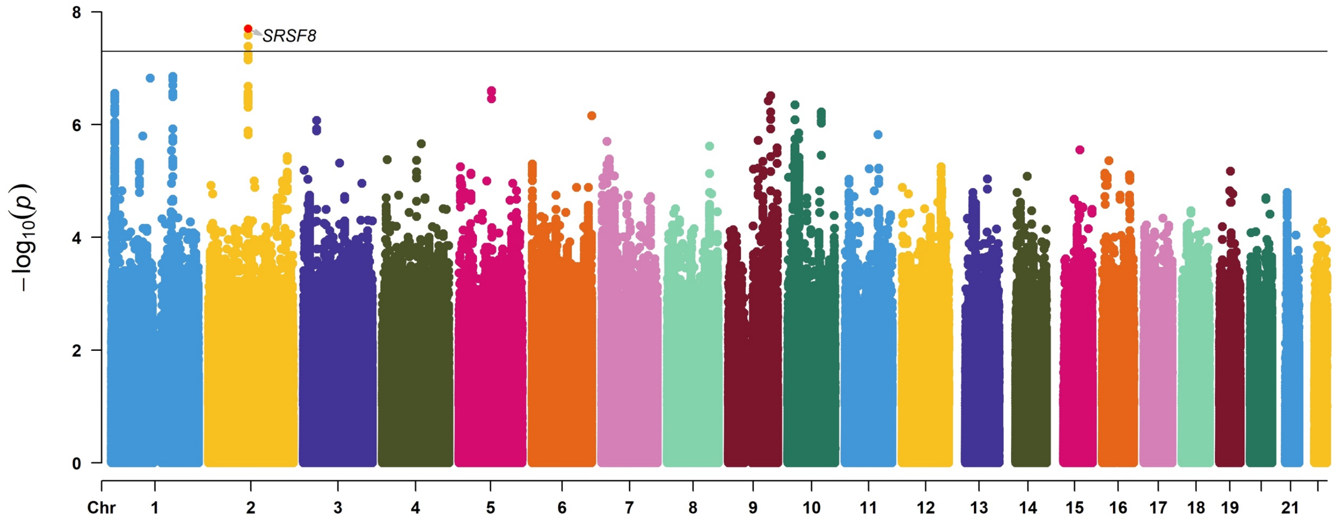


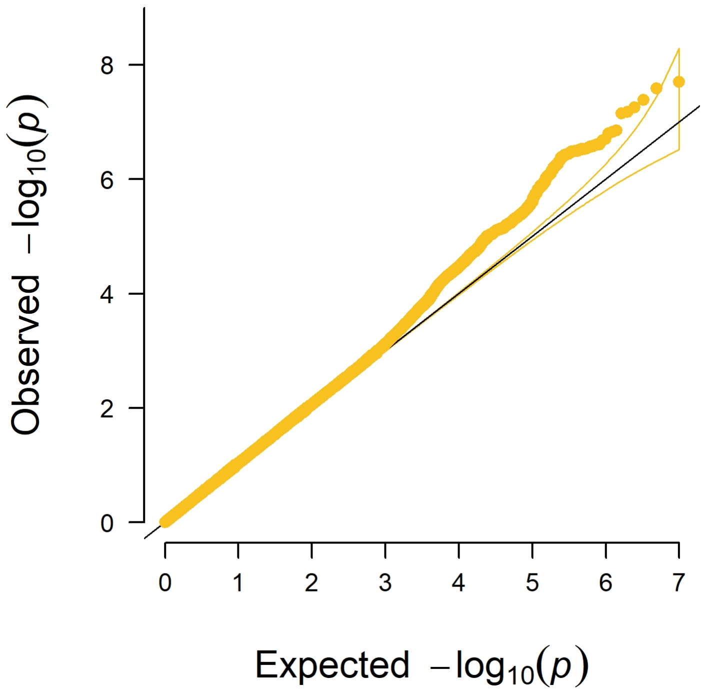


**Lambda gc = 1.046**

C. Louvain Modularity

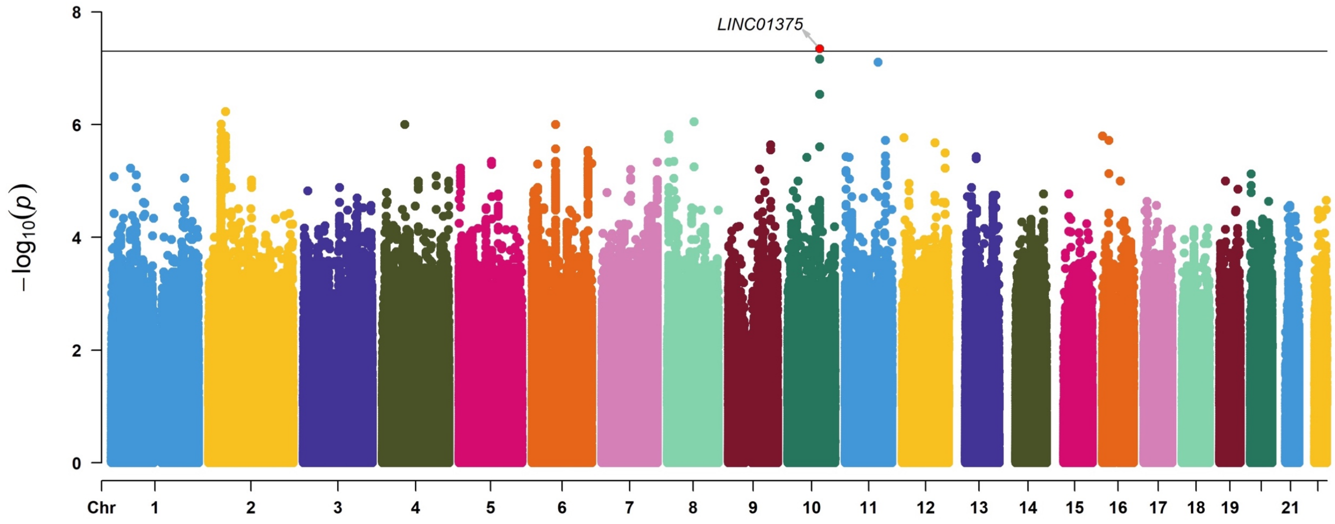


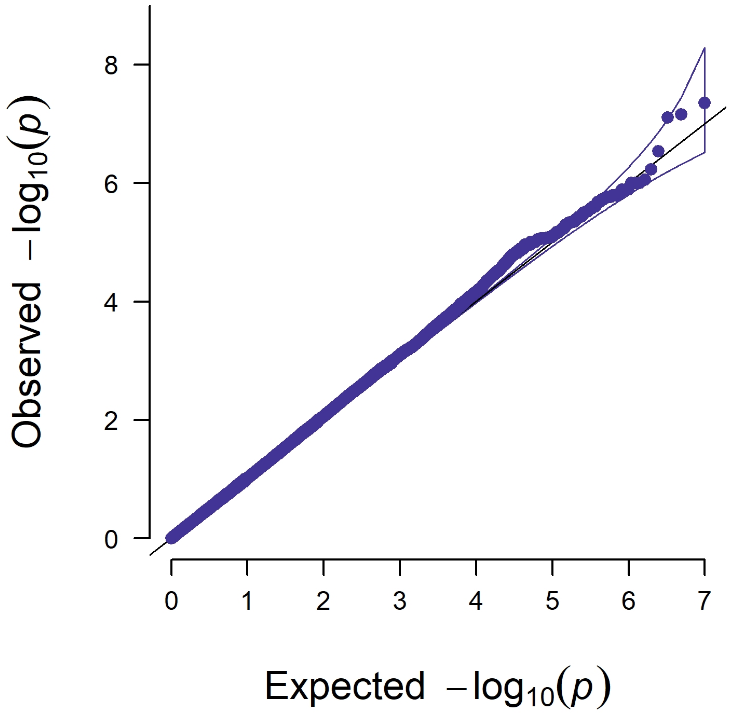


**Lambda gc = 1.034**

D. Transitivity

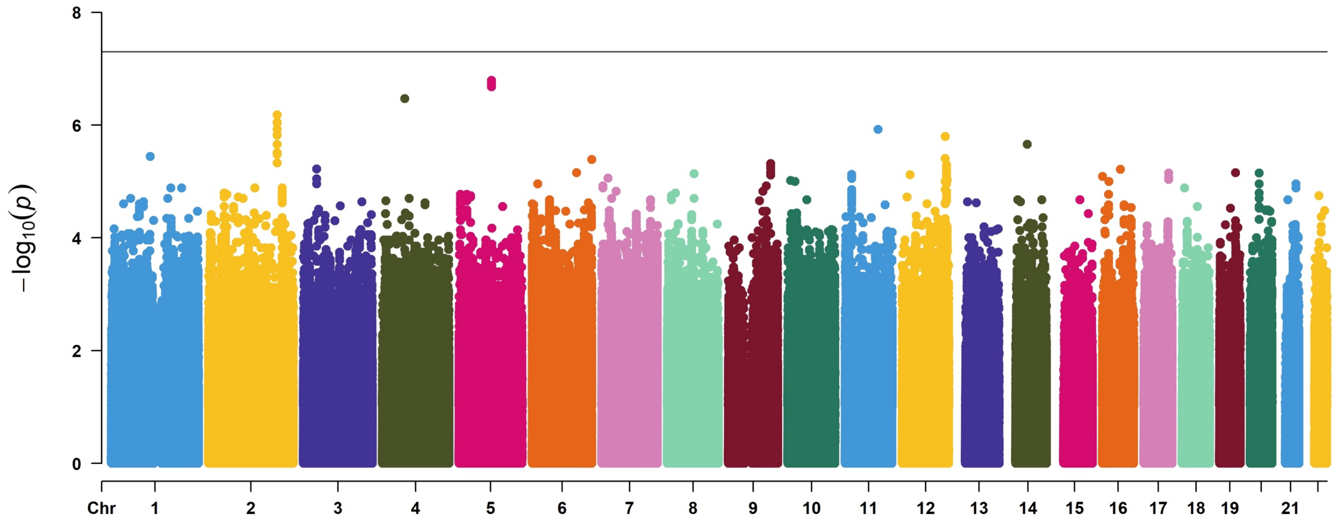


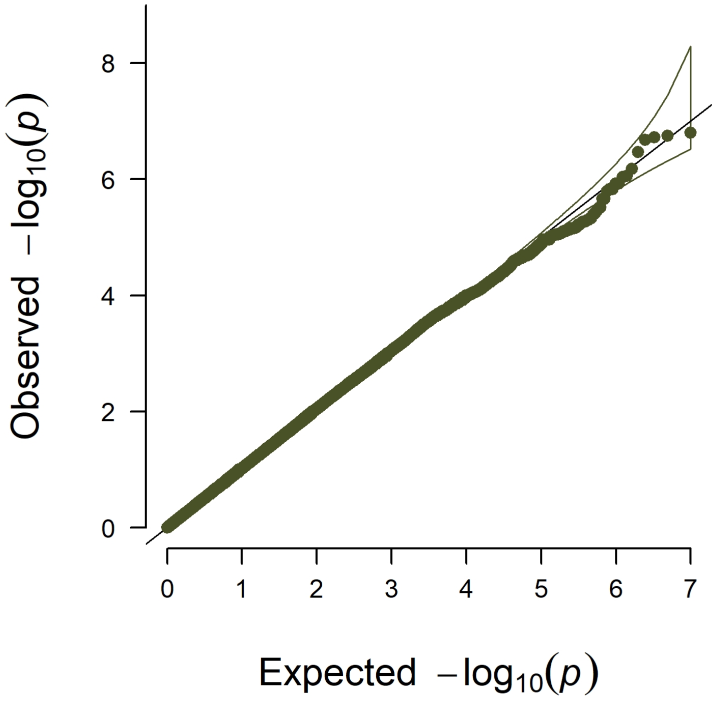


**Lambda gc = 1.027**

E. Local efficiency of default network

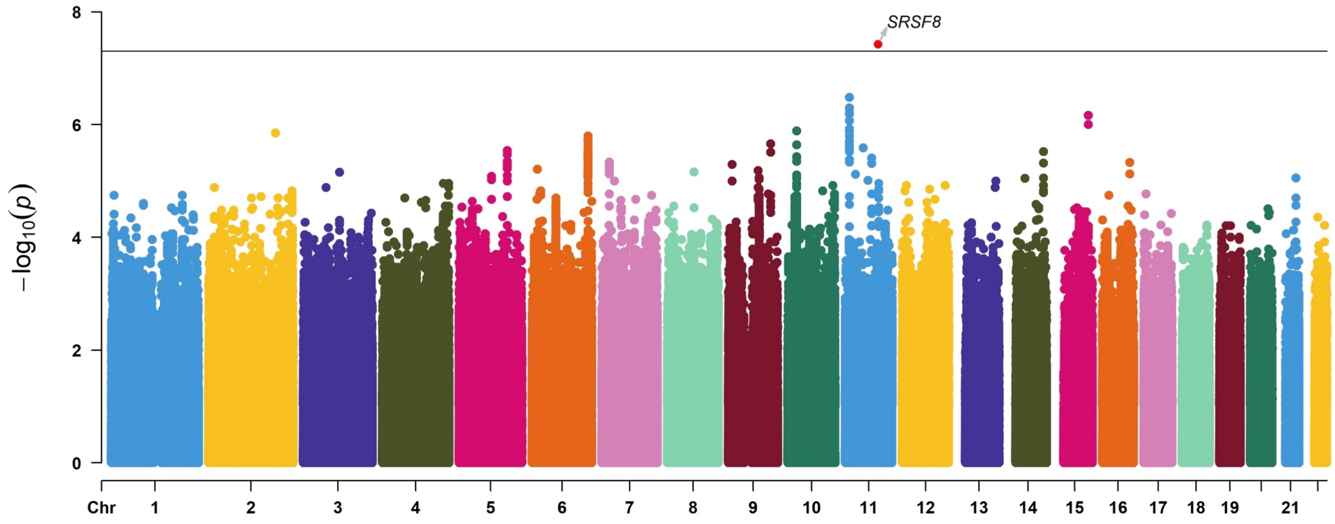


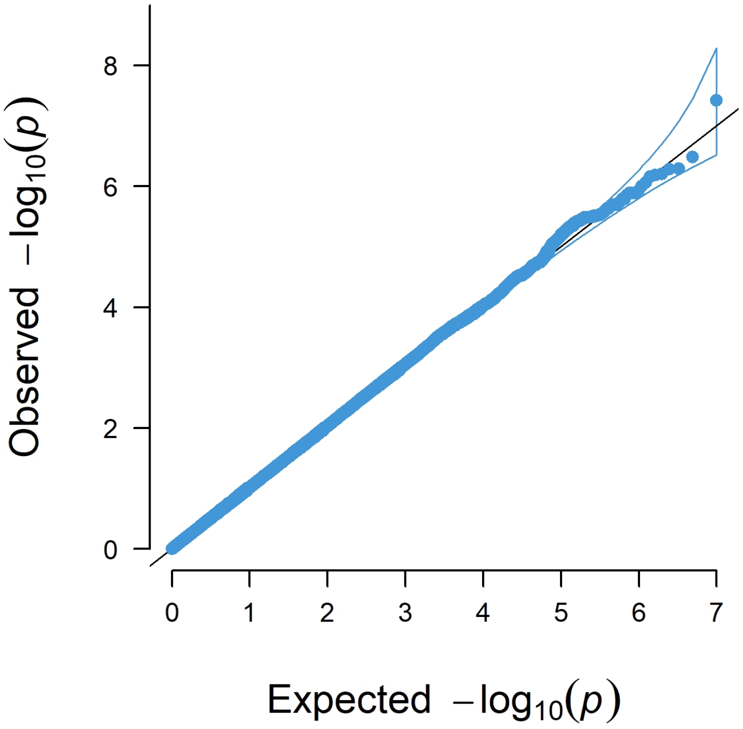


**Lambda gc = 1.024**

F. Local efficiency of dorsal attention network

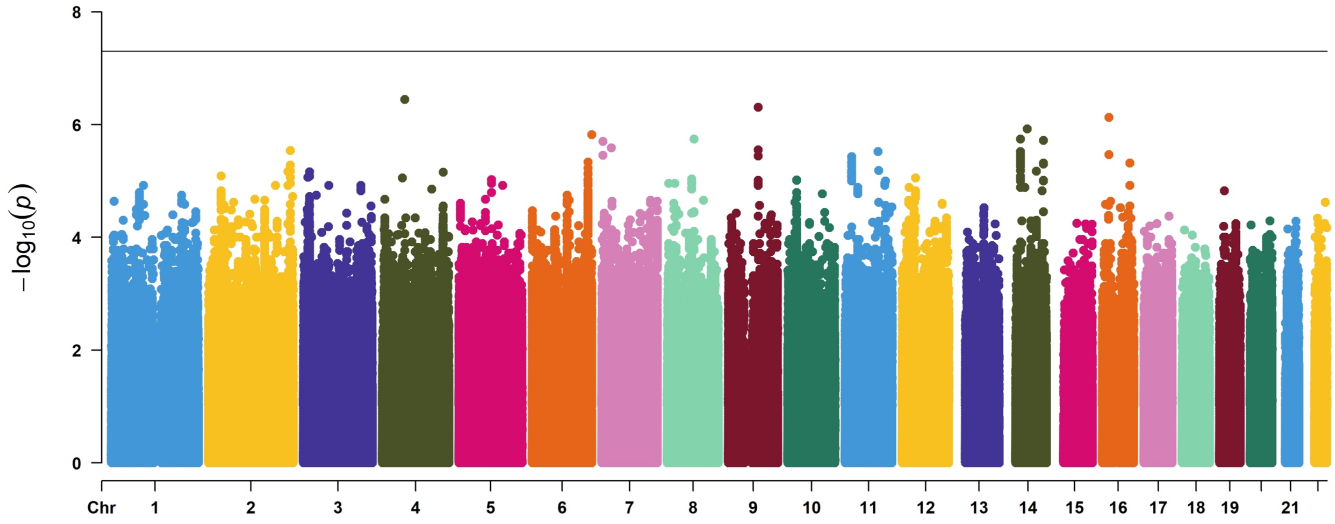


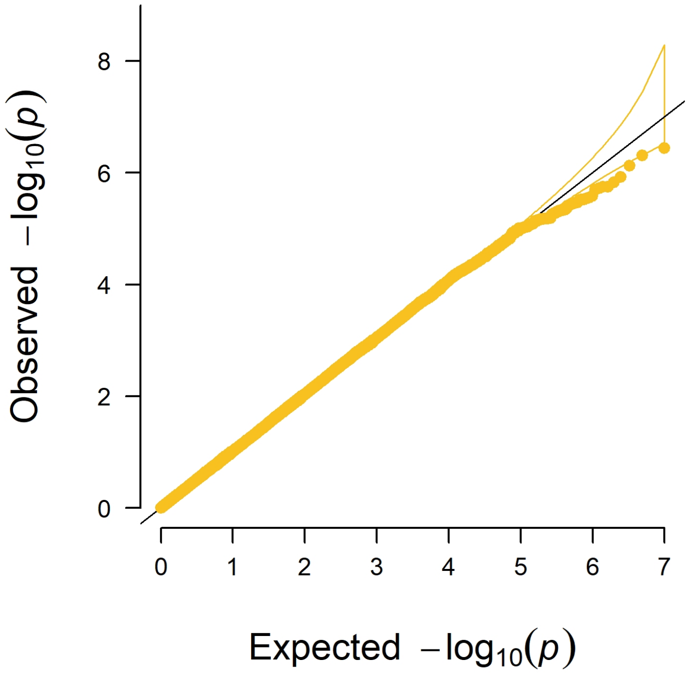


**Lambda gc = 1.021**

G. Local efficiency of frontoparietal network

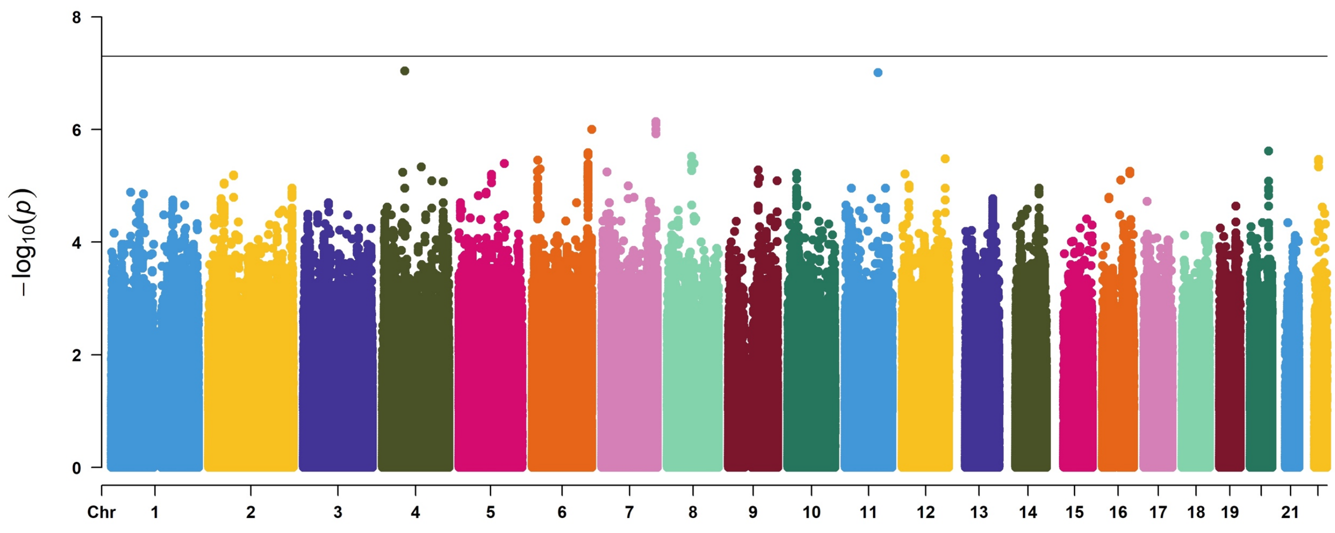


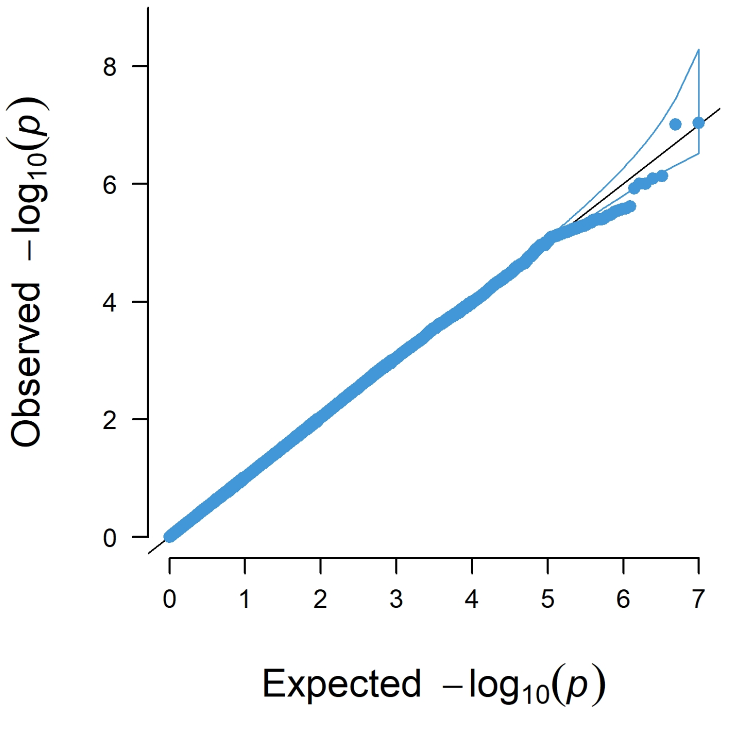


**Lambda gc = 1.020**

H. Local efficiency of limbic network

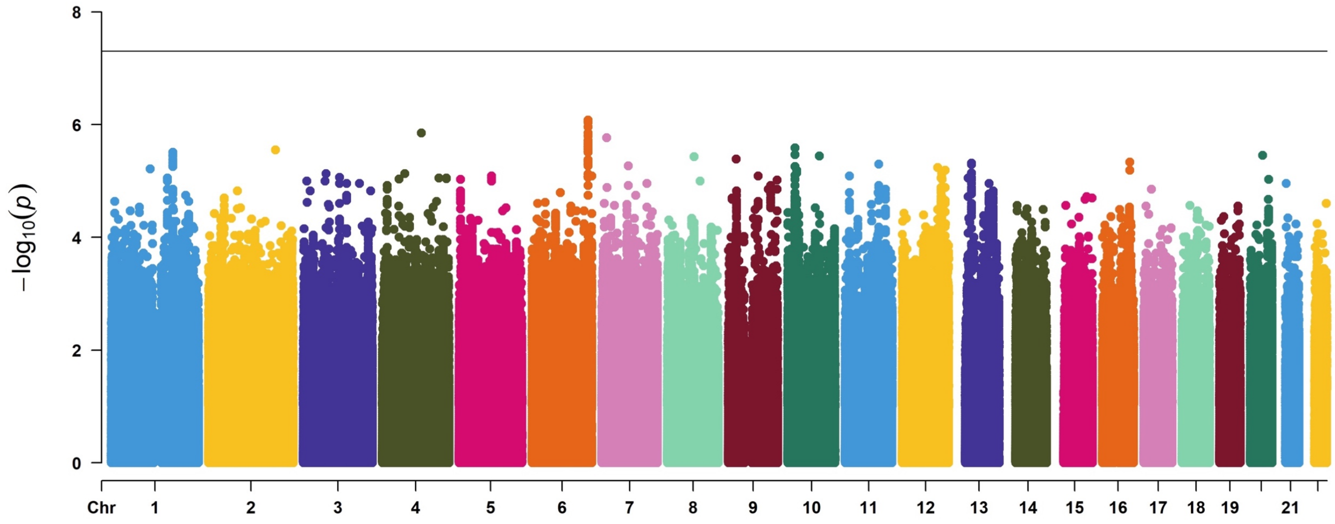


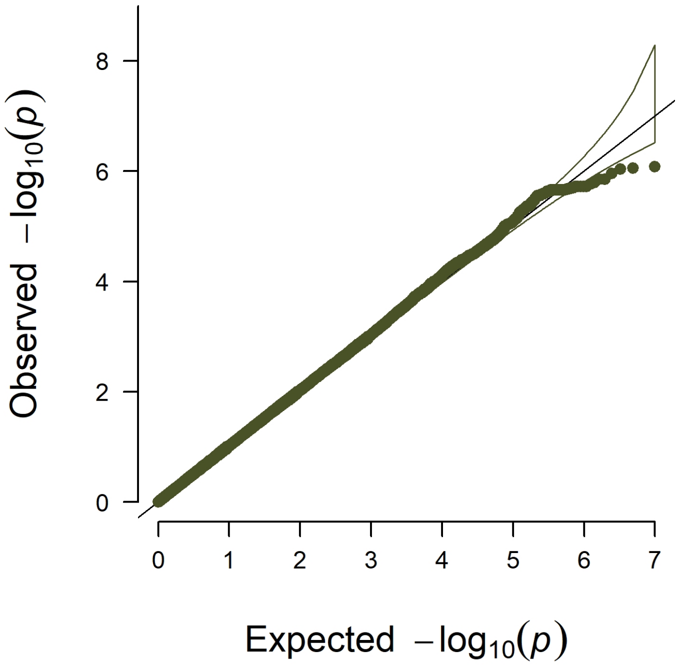


**Lambda gc = 1.022**

I. Local efficiency of salience network

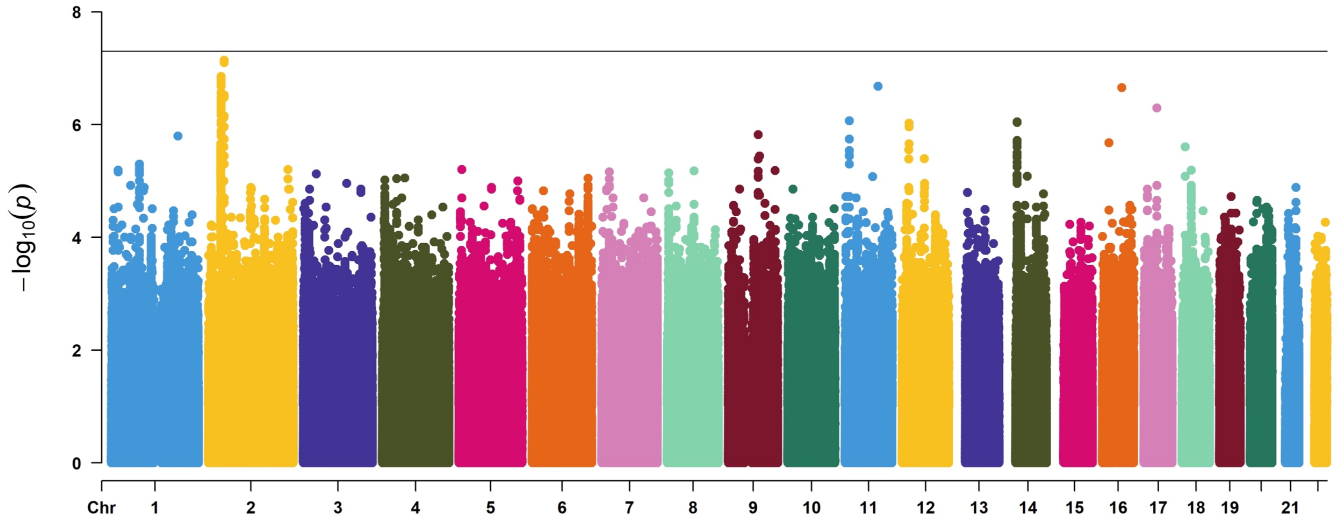


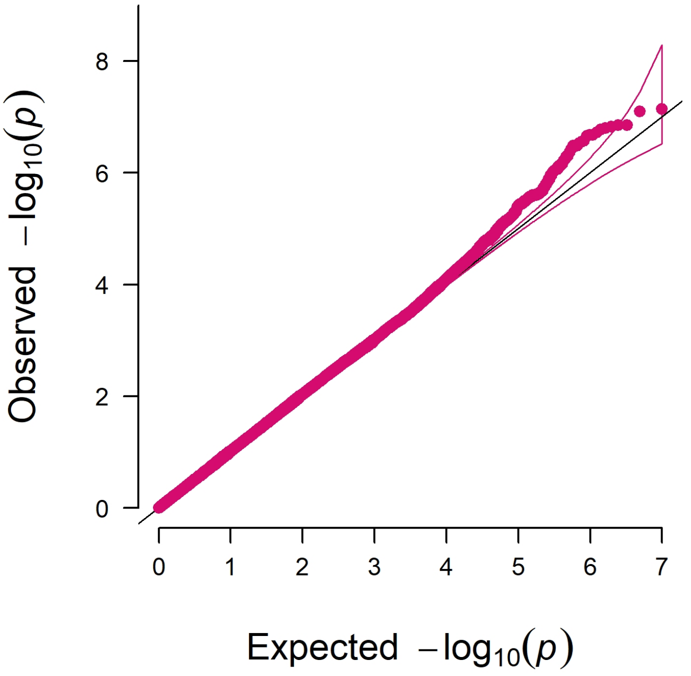


**Lambda gc = 1.019**

J. Local efficiency of somatomotor network

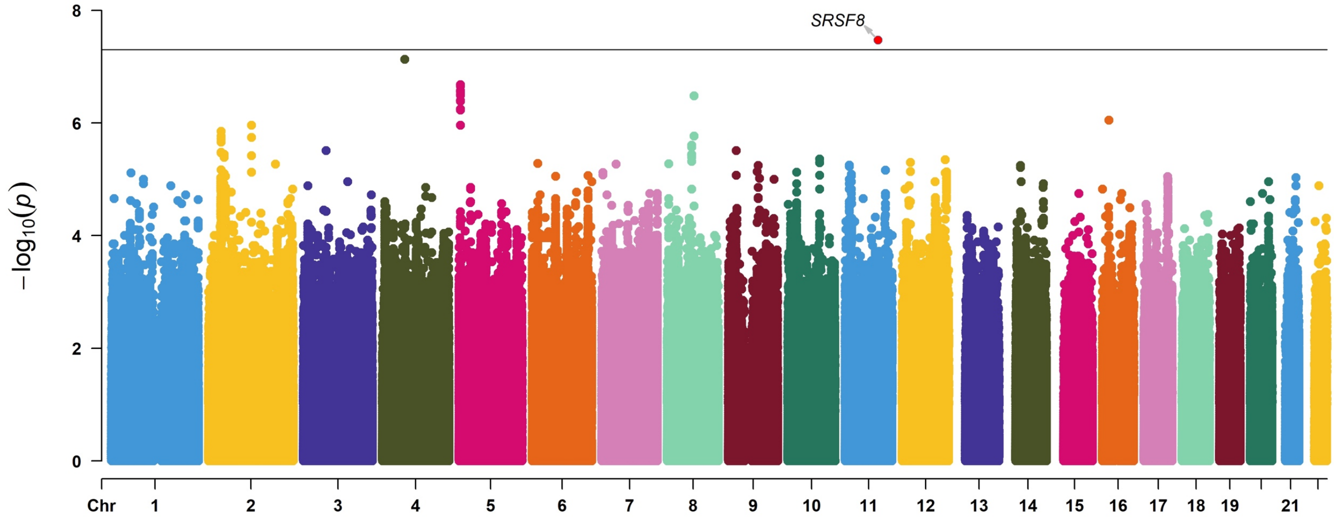


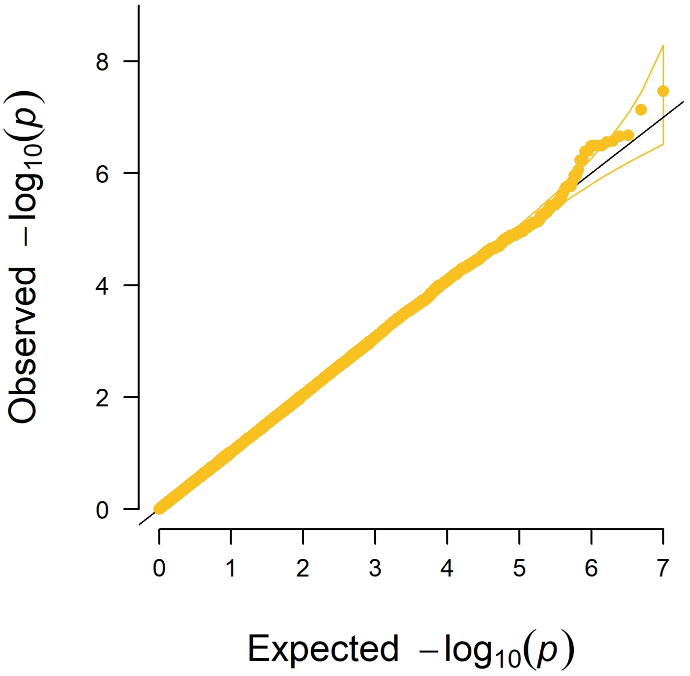


**Lambda gc = 1.025**

K. Local efficiency of visual network

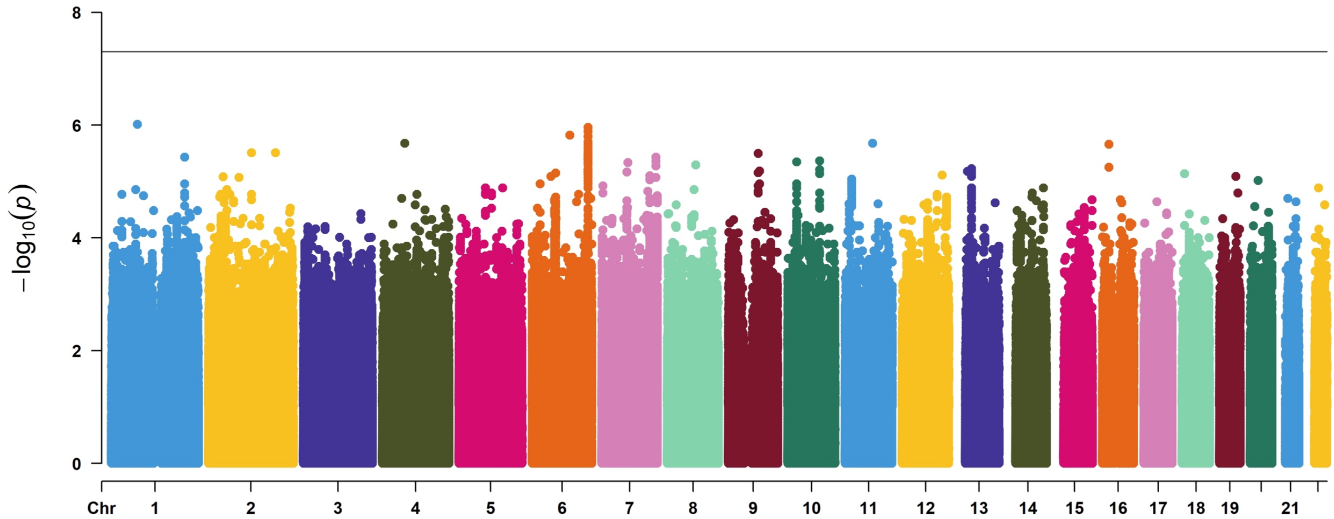


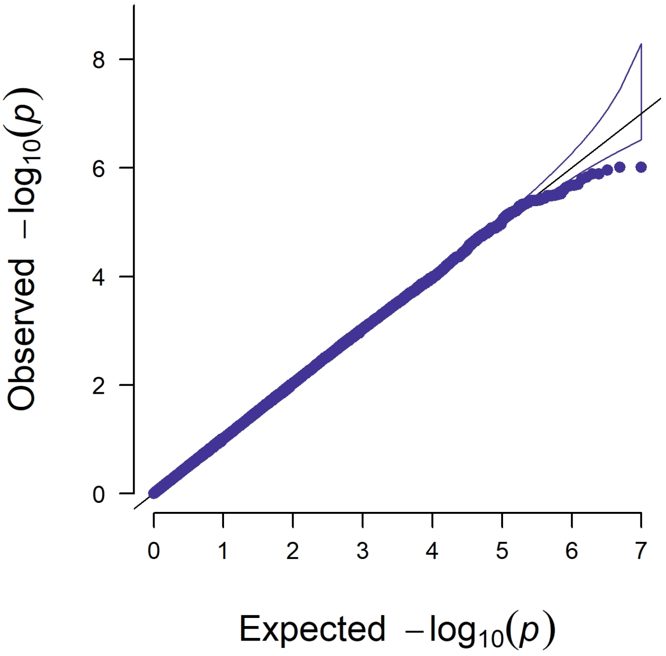


**Lambda gc = 1.018**

L. Strength of default network

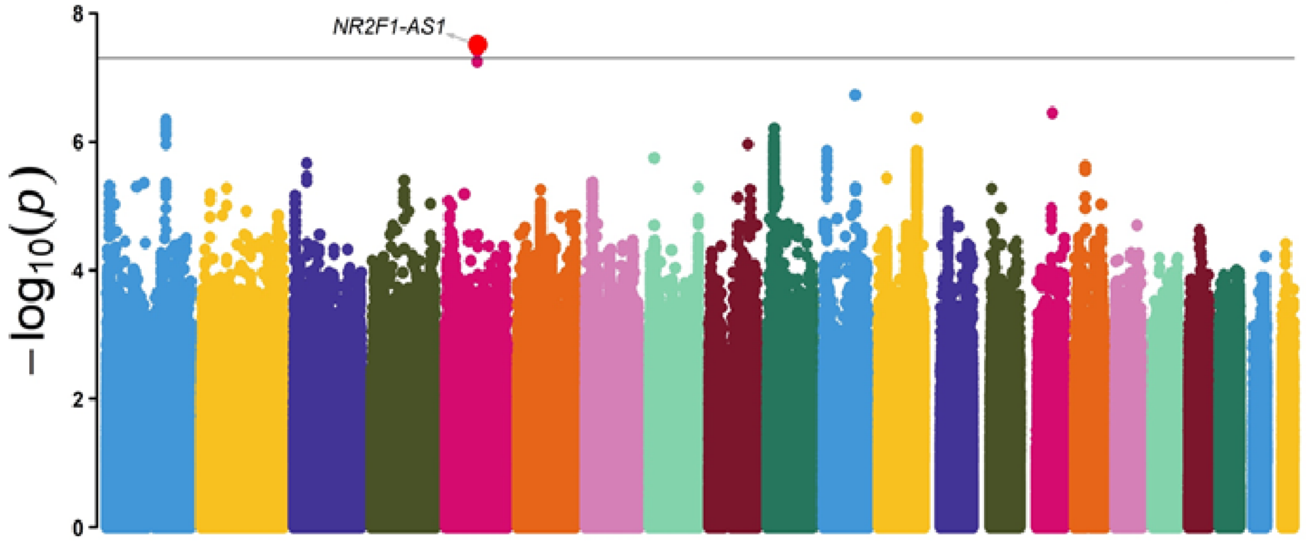


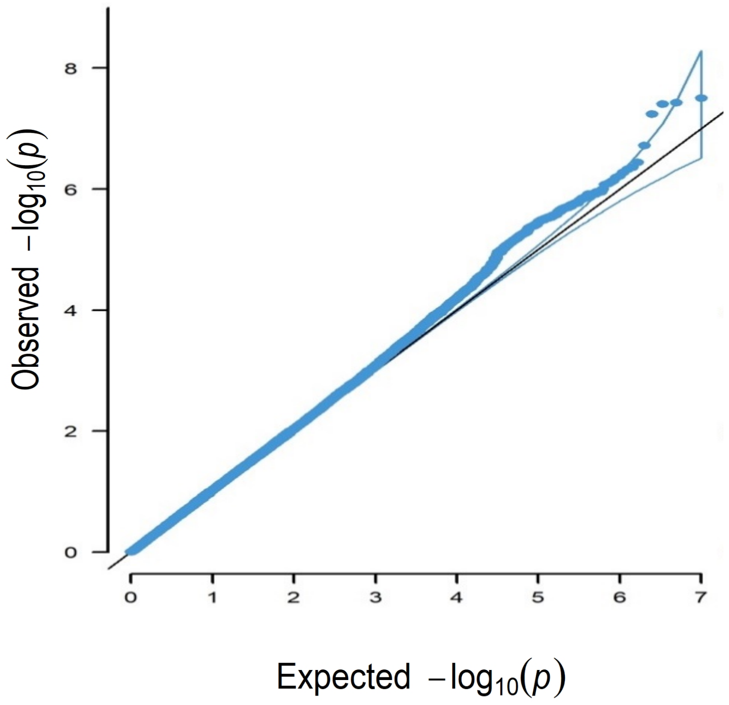


**Lambda gc = 1.026**

M. Strength of dorsal attention network

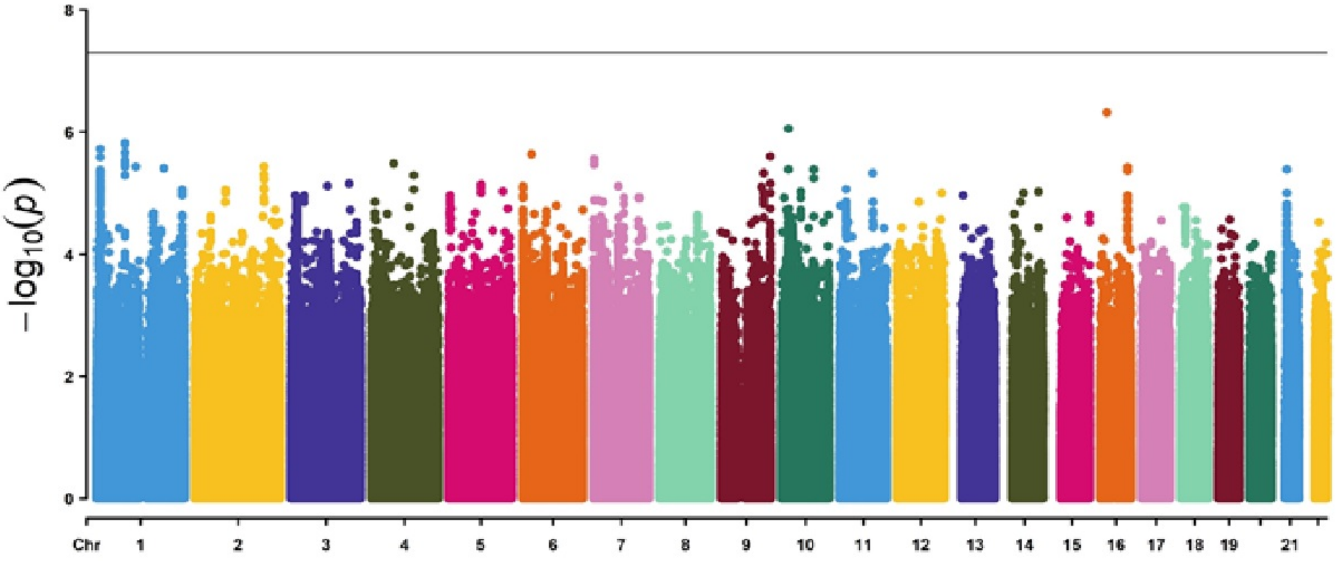


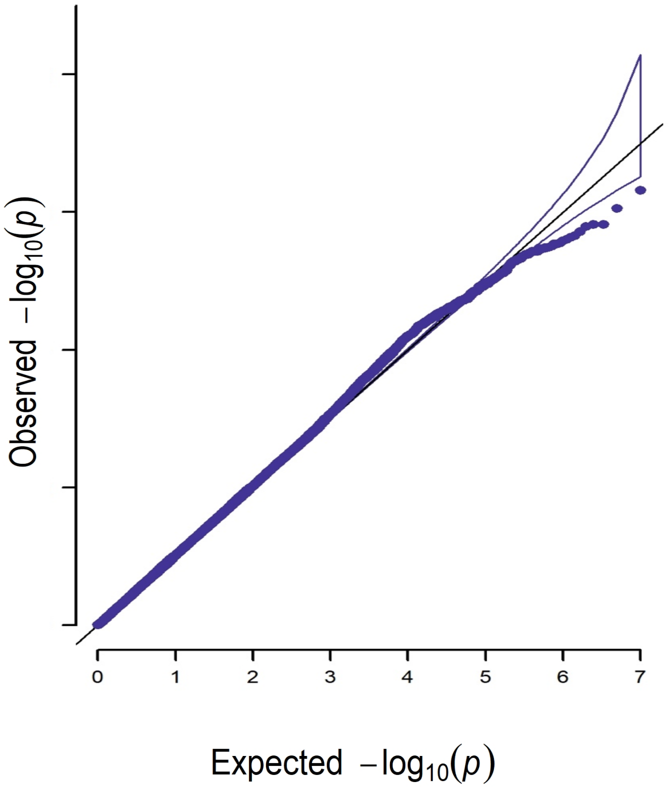


**Lambda gc = 1.033**

N.    Strength of frontoparietal network

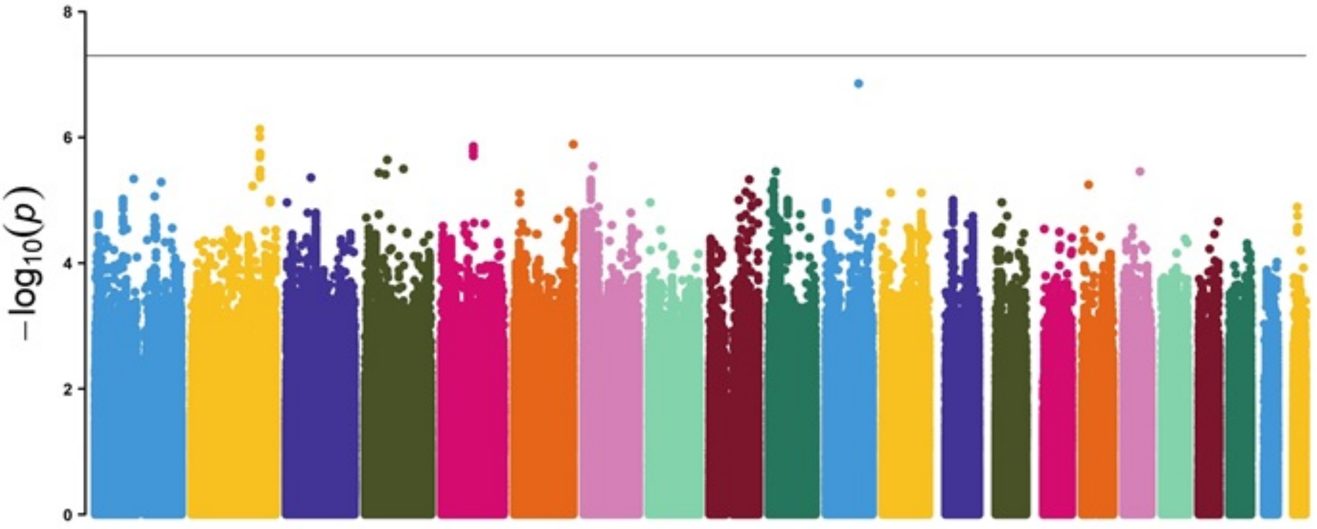


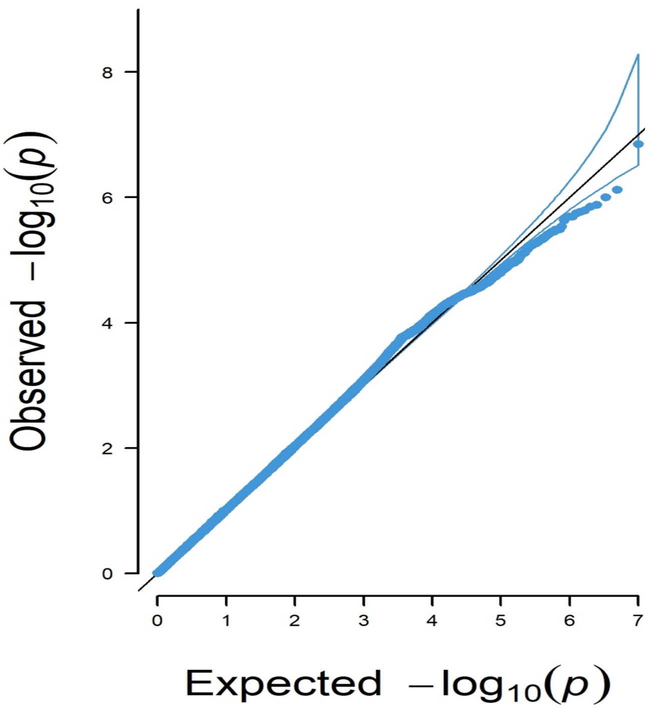


**Lambda gc = 1.041**

O. Strength of limbic network

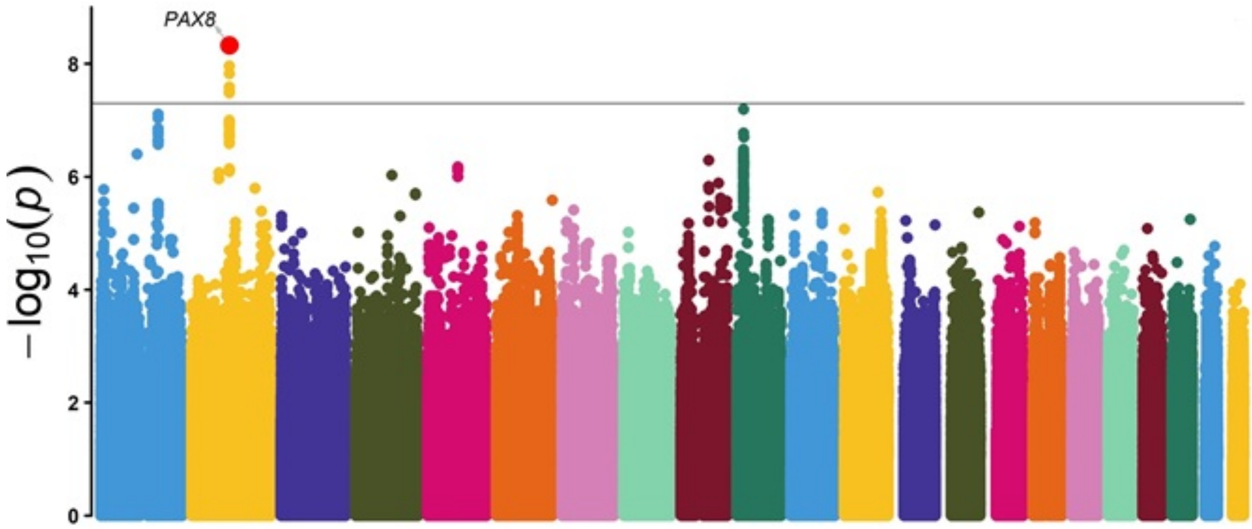


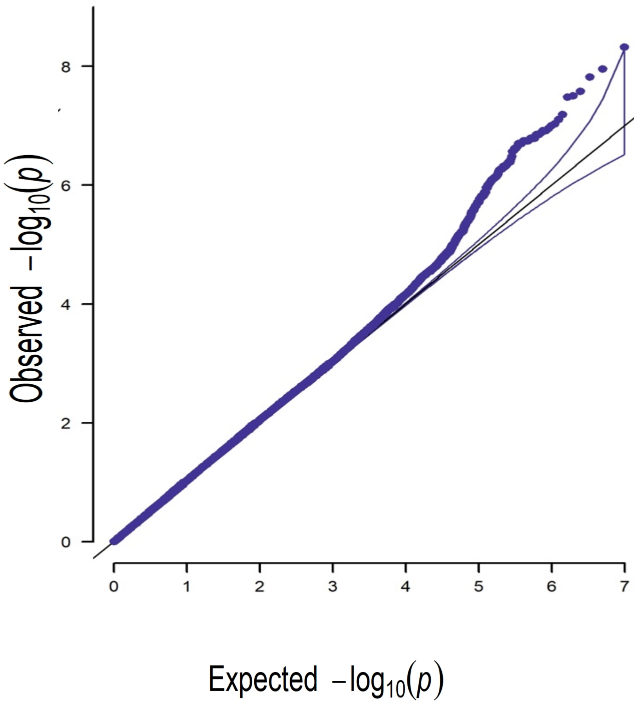


**Lambda gc = 1.013**

P. Strength of salience network

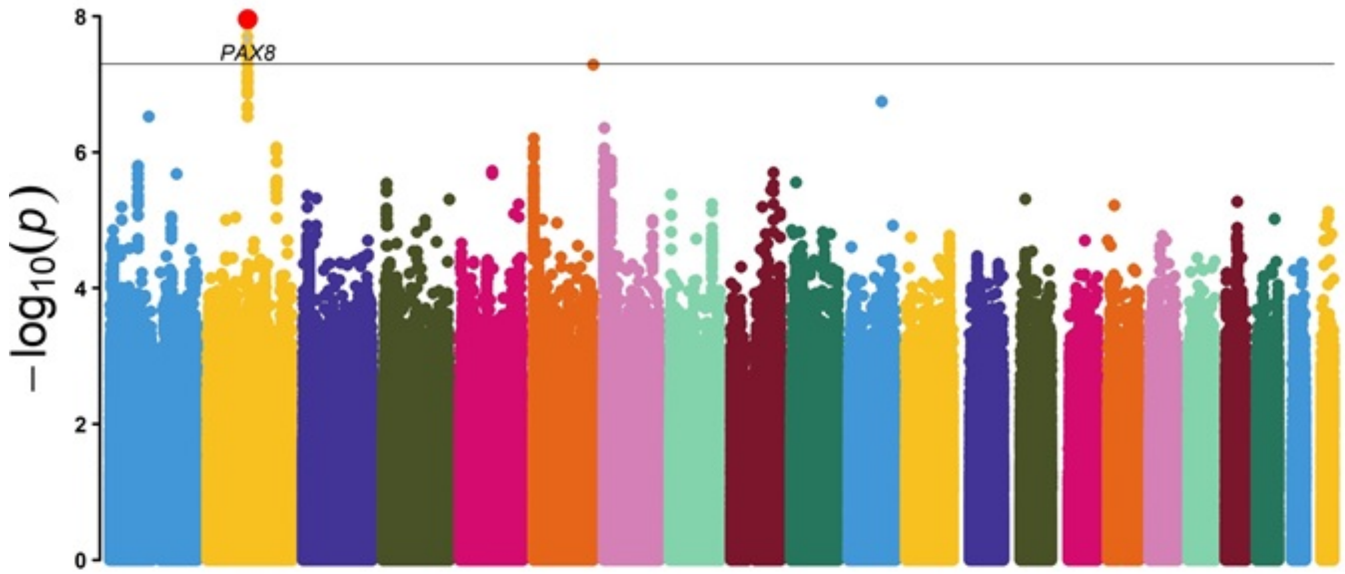


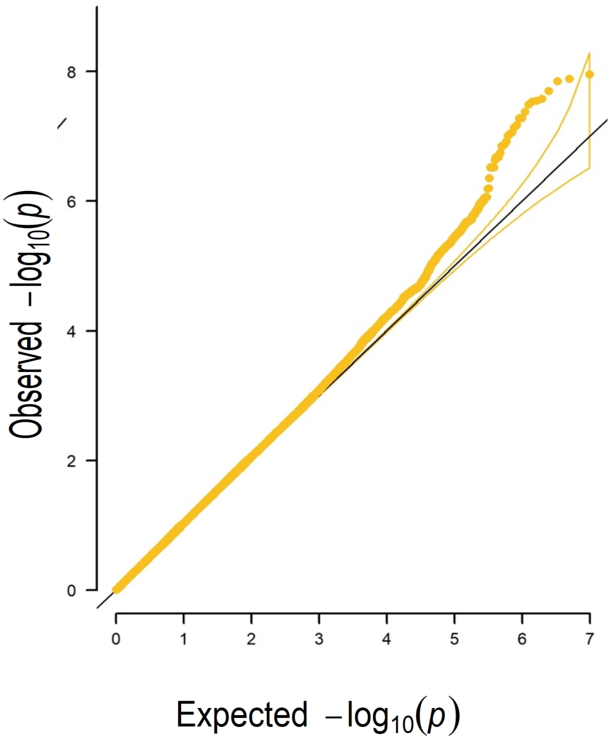


**Lambda gc = 1.022**

Q. Strength of somatomotor network

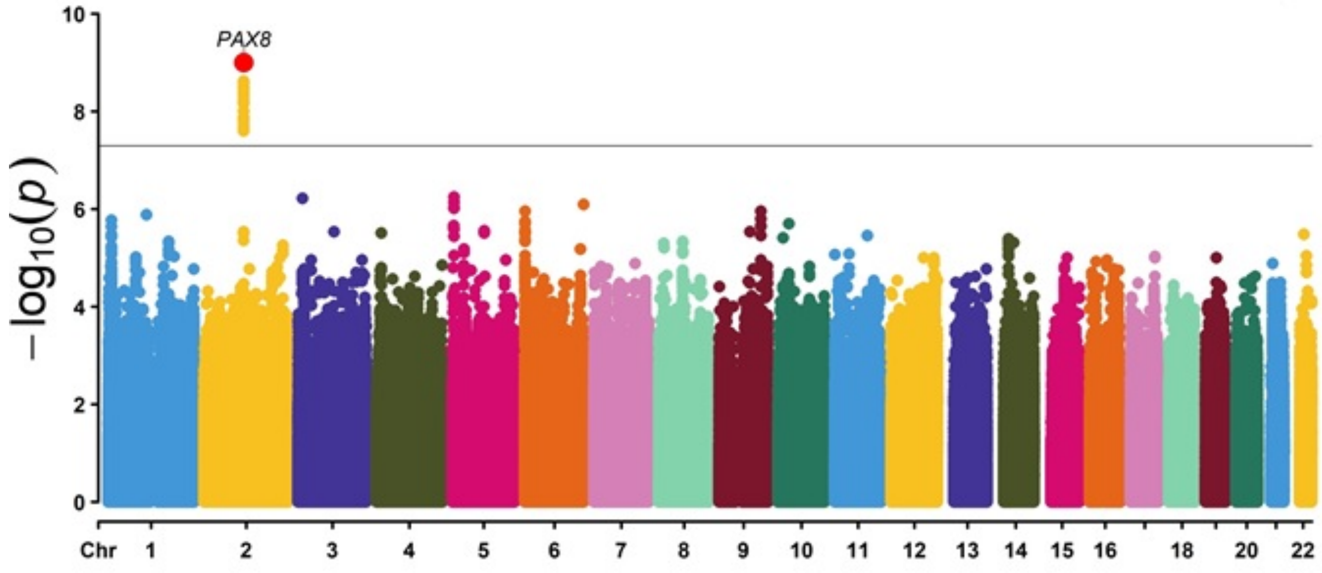


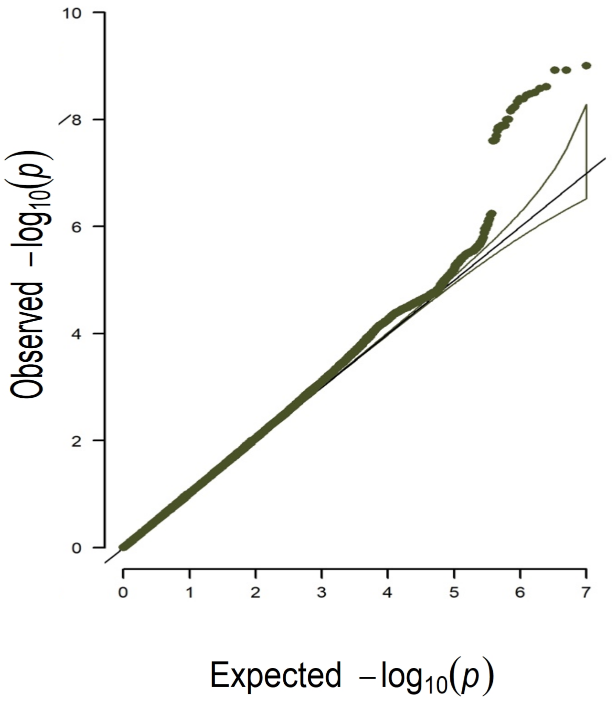


**Lambda gc = 1.037**

R. Strength of visual network

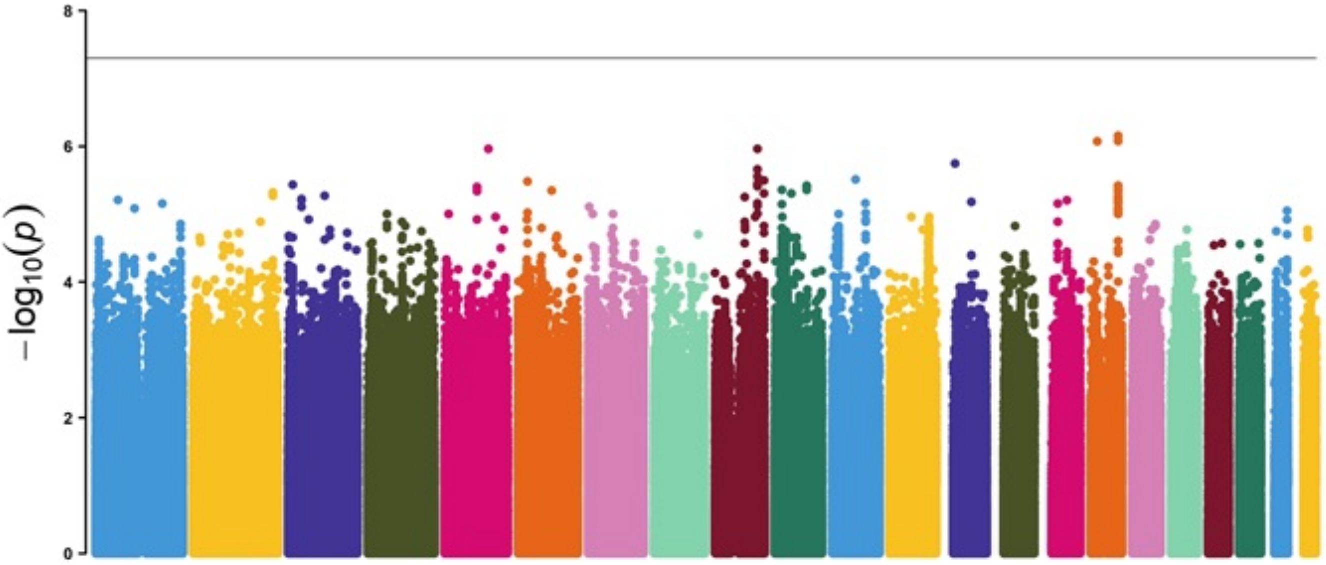


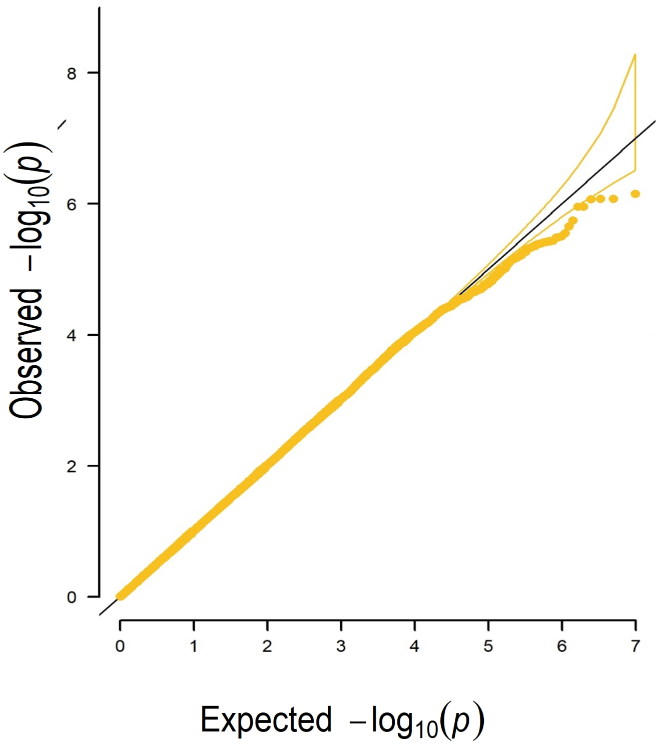


**Lambda gc = 1.034**

**Supplementary Fig. 3.** ***Heat map of gene expression levels across tissues for the list of genes found in the gene-based association analysis using FUMA Gene2Func.*** The colour bar represents the magnitude and direction of the gene expression – red represents higher expression of the genes compared to the cells filled in blue across tissue types.

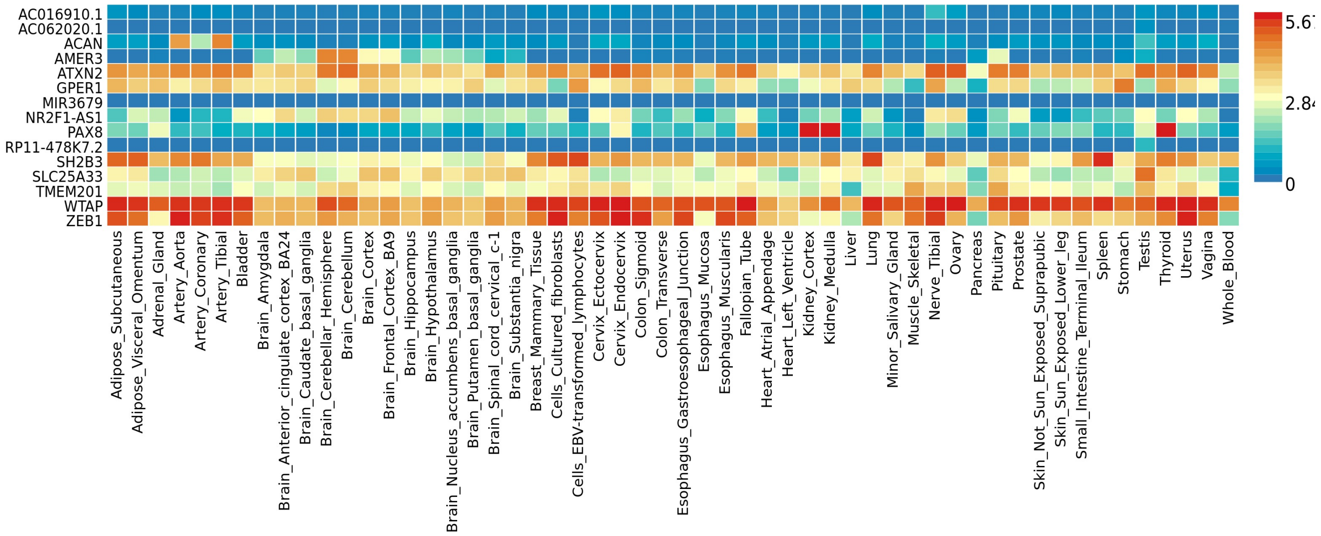
